# Phylogenetic background and habitat drive the genetic diversification of *Escherichia coli*

**DOI:** 10.1101/2020.02.12.945709

**Authors:** Marie Touchon, Amandine Perrin, Jorge André Moura de Sousa, Belinda Vangchhia, Samantha Burn, Claire L. O’Brien, Erick Denamur, David Gordon, Eduardo PC Rocha

**Affiliations:** Microbial Evolutionary Genomics, Institut Pasteur, CNRS, UMR3525, 25-28 rue Dr Roux, Paris, 75015, France; Ecology and Evolution, Research School of Biology, The Australian National University, 116 Daley Road, Acton, ACT, 2601, Australia; Department of Veterinary Microbiology, College of Veterinary Sciences & Animal Husbandry, Central Agricultural University, Selesih, Aizawl, Mizoram, 796014, India; School of Medicine, University of Wollongong, Northfields Ave Wollongong, NSW 2522, Australia; Université de Paris, IAME, UMR 1137, INSERM, 75018, Paris, France; AP-HP, Laboratoire de Génétique Moléculaire, Hôpital Bichat, 75018, Paris, France; Sorbonne Université, Collège doctoral, F-75005 Paris, France

**Keywords:** local adaptation, gene repertoire, mobile genetic elements, horizontal gene transfer, freshwater isolates

## Abstract

*Escherichia coli* is a commensal of birds and mammals, including humans. It can act as an opportunistic pathogen and is also found in water and sediments. Since most population studies have focused on clinical isolates, we studied the phylogeny, genetic diversification, and habitat-association of 1,294 isolates representative of the phylogenetic diversity of more than 5,000, mostly non-clinical, isolates originating from humans, poultry, wild animals and water sampled from the Australian continent. These strains represent the species diversity and show large variations in gene repertoires within sequence types. Recent gene transfer is driven by mobile elements and determined by habitat sharing and by phylogroup membership, suggesting that gene flow reinforces the association of certain genetic backgrounds with specific habitats. The phylogroups with smallest genomes had the highest rates of gene repertoire diversification and fewer but more diverse mobile genetic elements, suggesting that smaller genomes are associated with higher, not lower, turnover of genetic information. Many of these small genomes were in freshwater isolates suggesting that some lineages are specifically adapted to this environment. Altogether, these data contribute to explain why epidemiological clones tend to emerge from specific phylogenetic groups in the presence of pervasive horizontal gene transfer across the species.

## Introduction

The integration of epidemiology and genomics has greatly contributed to our understanding of the population genetics of epidemic clones of pathogenic bacteria. However, the forces driving the emergence of these lineages in species where most clades are dominated by commensal or environmental strains remain unclear. *Escherichia coli* is a commensal of the gut microbiota of mammals and birds (primary habitat)^1–3^, and has been found in host-independent secondary habitats including soil, sediments, and water^4–7^. Yet, some *E. coli* strains produce virulence factors endowing them with the ability to cause a broad range of intestinal or extra-intestinal diseases (pathotypes) in humans and domestic animals^8–13^. Many of these are becoming resistant to multiple antibiotics at a worrisome pace^14, 15^.

Studies on *E. coli* were seminal in the development of bacterial population genetics^16^. They showed moderate levels of recombination in the species^3, 17–19^, and a strong phylogenetic structure with eight main phylogroups, among which four (A, B1, B2 and D) represent the majority of the strains and four others (C, E, F and G) are rarer^20–22^. Strains differ in their phenotypic and genotypic characteristics within and across phylogroups^2, 3, 23, 24^, and their isolation frequency depends on factors such as host species, diet, sex, age^25–27^, body mass^28^, but also climate^29, 30^, and geographic location^31^. Strains of phylogroups A and B1 appear to be more generalists since they can be isolated from all vertebrates^2^ and are often isolated from secondary habitats^7, 32–35^. *E. coli* strains able to survive and persist in water environments usually belong to the B1 phylogroup^7, 33, 34^. In contrast, the extraintestinal pathogenic strains usually belong to phylogroups B2 and D^36–38^. Genome size also differs among phylogroups, with A and B1 strains having smaller genomes than B2 or D strains^23^. The phylogenetic vicinity of geographically remote *E. coli* isolates, and the co-isolation of phylogenetically distant strains, supports the hypothesis that strains circulate rapidly^39, 40^. The genome of the species is also remarkably plastic, since only about half of the average genome is present across most strains of the species (core or persistent genome) and the pan-genome vastly exceeds the size of the typical genome^41–44^. Interestingly, the rapid circulation of strains and the high plasticity of their genomes have not erased the associations of certain clades with certain isolation sources. In consequence, such associations might reflect local adaptation^16, 45^, which would suggest frequent genetic interactions between the novel adaptive changes and the strains’ genomic background.

Understanding how the evolution of gene repertoires is shaped by population structure and habitats requires large-scale comparative genomics of samples with diverse sources of isolation representative of natural populations of *E. coli*. Most of the efforts of genome sequencing have been devoted to study pathogenic lineages and very few genomic data are available for commensal strains, especially in wild animals, and environmental strains. Here, we analysed the genomes of a large collection of *E. coli* strains collected across many human, domestic and wild animal and environmental sources in different geographic locations from the Australian continent. This collection is dominated by non-clinical isolates, corresponding to the main habitats of the species. We sought to understand the dynamics of the evolution of gene repertoires and how it was driven by mobile genetic elements. The analysis of the isolation sources in the light of phylogenetic structure and genome variation suggests that adaptation varies with the habitat and the phylogenomic background. This contributes to explain why known epidemiological clones of the species emerge from specific phylogenetic groups, even though virulence strongly depends on the acquisition of virulence factors by horizontal gene transfer.

## Results

### Very rapid initial divergence of gene repertoires becomes linear with time

We sequenced and annotated the genomes of 1,294 *E. coli sensu stricto* strains selected from more than 3,300 non-human vertebrate hosts, 1,000 humans and 800 environmental samples between 1993 and 2015, chosen to represent the phylogenetic diversity of the species (Materials and Methods, Fig. 1a, Supplementary Notes). All samples were collected by a single team, spanning a 20 year-period, from different regions in a single isolated continent (Australia). The origin of each strain was accurately characterized and the genomes were uniformly annotated and analyzed using the same bioinformatics processes. The strains were isolated from humans, domesticated and wild animals, representing the primary habitat of *E. coli*, and from freshwater, representing its secondary habitat^3^. Less than 22% of the samples were recovered from clinical situations. A series of controls confirmed that the sequences were of high quality and contained the known essential genes (Supplementary Notes). The genomes varied widely in size from 4.2 to 6.0 Mb (average 5 Mb), but had similar densities of protein-coding sequences (∼87%) and GC content (50.6%, Supplementary Fig. 1 and Supplementary Table 1).

**Fig. 1:**
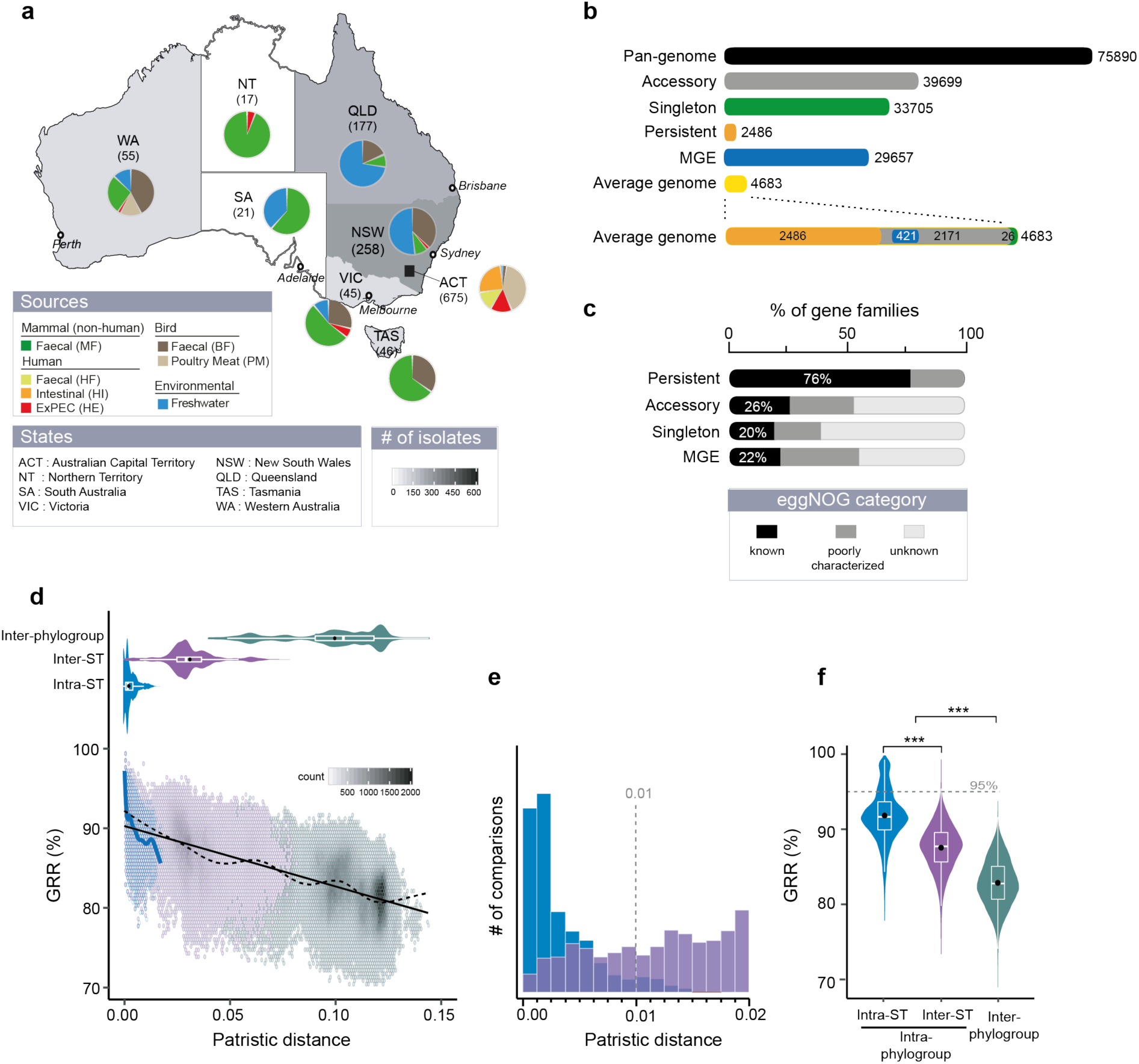
The genetic diversity of Australian *E. coli*. **(a)** Distribution of isolates per region and per source. **(b)** The pan-genome is composed of 75,890 gene families, of which 33,705 are singletons (in green, present in a single genome), 2,486 persistent (in gold, present in at least 99% of genomes), the remaining being accessory (in grey). 29,657 gene families (39% of the pan-genome) were related to mobile genetic elements (MGE). **(c)** Percentage of the different EggNOG categories (see insert) in the persistent, accessory and singleton gene families and among genes associated to MGE. **(d)** [Top] Violin plots of the patristic distance computed between pairs of genomes. [Bottom] Association between GRR (Gene Repertoire Relatedness) and the patristic distance across pairs of genomes. Due to the large number of comparisons (points), we divided the plot area in regular hexagons. Color intensity is proportional to the number of cases (count) in each hexagon. The linear fit (black solid line, linear model (lm)) was computed for the entire dataset (1,294 genomes, Y=90.2-75.7*X, R^2^=0.49, P<10^-4^). The spline fit (generalized additive model (gam)) was computed for the whole (in black dashed line) or the intra-ST (in blue solid line) comparisons. There was a significant negative correlation between GRR and the patristic distance (Spearman’s rho = −0.67, P<10^-4^). **(e)** Histograms of the number of intra-ST (in blue) and inter-ST (in purple) comparisons at short evolutionary scales. **(f)** Violin plots of the intra-ST, inter-ST and inter-phylogroup GRR (%). (d-e-f) All the distributions were significantly different (Wilcoxon test, P<10^-4^), the same color code was used and described in (d).

The pan-genome contained 75,890 gene families that were classified as *persistent* (3%, gene families present in ≥ 99% of the genomes), *singletons* (44%, present in a single genome), or *accessory* (the remaining) (Fig. 1b, Supplementary Fig. 2). The persistent gene families are a tiny fraction of the pan-genome, but account for half of the average genome. They were used to build a robust phylogeny of the species, which was rooted using genomes from other species in the genus (Supplementary Fig. 3). In contrast, singletons are almost half of the gene families of the pan-genome, but less than 1% of the average genome. As a consequence, the pan-genome is open, as measured by the fit to a Heaps’ law model^46^, and increases on average by ∼26 protein coding genes with the inclusion of a new genome (Supplementary Fig. 2). Singletons are smaller than the other genes and tend to be located at the edge of contigs (44%). Hence, some of these singletons may result from sequencing and assembly artifacts (Supplementary Notes and Supplementary Fig. 4). When all the singletons were excluded, the pan-genome still remained open (Supplementary Fig. 2). Most singletons (80%) and accessory (74%) gene families, but also a surprisingly high number of persistent gene families (24%), lacked a clear functional assignment as given by the EggNOG database^47^ (Fig. 1c). Hence, we are still ignorant of the function, or even the existence, of many genes of the species.

Traditional epidemiological studies of *E. coli* focused on multilocus sequence types (ST) and/or the O- and H-serotypes (often the O:H combination). These epidemiological units regroup strains in terms of sequence similarity in a few persistent genes (ST) or in key traits related to the cell envelope (the LPS structure and the flagellum). However, it is unclear if these types systematically regroup strains with similar gene repertoires. We identified 442 distinct STs, of which 61% are represented by a single strain. A few STs are very abundant in our dataset: 20 include more than 10 genomes each and encompass 40% of the dataset. The intra-ST genetic distances are 10-times smaller than distances between other pairs of genomes (0.003 vs. 0.03, Fig. 1d). Yet, 6% of intra-ST comparisons have more than 0.01 substitutions per position showing extensive genetic diversity at the genome level (Fig. 1e). Some O-groups are abundant, e.g., O8, O2 and O1 (each present in >50 genomes) but almost half of the groups occur in a single genome and 43% of the strains could not be assigned an O-group (even when the *wzm*/*wzt* and *wzx*/*wzy* genes were present). In contrast, most H-types were previously known (87%). We found 311 combinations of O:H serotypes among the 726 typeable genomes. Of these, 64% are present in only one genome,17% are in multiple STs and 7% in multiple phylogroups (e.g. O8:H10). Conversely, half of the 95 STs with more than one genome have multiple O:H combinations, e.g. ST10 has 24. These results confirm that surface antigens and their combinations change quickly and are homoplasic. They also show extensive variation of gene repertoires within STs. The gene repertoire relatedness (GRR) between genomes (see Methods) decreases very rapidly with phylogenetic distance for closely related strains, as revealed by spline fits (Fig. 1d). Similar results were observed when removing singletons, which only account for on average 0.5% of the genes in genomes, suggesting that this result is not due to annotation or sequencing errors (Supplementary Fig. 6). As a consequence, 85% of the intra-ST comparisons have a GRR lower than 95% (corresponding to ∼235 gene differences per genome pair), and some as little as 77% (Fig. 1f). Hence, even genomes of the same ST can differ substantially in the sequence of other persistent genes and in the overall gene repertoires.

To check if the dataset is representative of the species and can be used to assess its diversity, we compared it with the ECOR collection^48^ and the complete genomes available in RefSeq (Materials). All datasets had similar nucleotide diversity (Supplementary Fig. 5a and Supplementary Table 1). Using rarefied datasets, to compare sets of same size, ours had the largest pan-genome, partly because of a larger number of singletons (Supplementary Fig. 5b-d). Our dataset also had the highest *α*-diversity for the three typing schemes (STs, O-, H-serotypes, Supplementary Table 1). Since the gene repertoire diversity of *E. coli* in Australia is at least as high as that of ECOR and RefSeq, we studied the variation in gene repertoires beyond the intra-ST level. After the rapid initial drop in GRR described above, the values of this variable decrease linearly with phylogenetic distances (Fig. 1d). The average values of GRR given by the regression vary between 90% for very close genomes and 80% for the most distant ones. The variance around the regression line is constant and a spline fit shows few deviations around the regression line. This is consistent with a model where initial divergence in gene repertoires is driven by rapid turnover of novel genes. After this initial process, divergence in gene repertoires increases linearly with patristic distance.

### Phylogroups vary in the rates of gene repertoire diversification

We used the species phylogeny to study the associations between phylogroups and genetic diversity (Fig. 2a). The tree showed seven main phylogenetic groups very clearly separated by nodes with 100% bootstrap support. The 17 phylogroup C strains were all included within the B1 phylogroup and were thus grouped with the latter in this study. For the rest, the analysis showed a good correspondence between the assignment into the known phylogroups - A, B1, B2, D, E, F, and G – and the different clades of this tree. In line with the literature^40^, four major phylogroups were very abundant - A (24% of the dataset), B1 (24%), B2 (25%) and D (14%) – whereas the others were rarer. The nucleotide diversity of the phylogroups is very dependent on their phylogenetic structure, since some clades have more closely related clusters of strains than others (Supplementary Fig. 7). Nevertheless, nucleotide diversity, patristic distances, and Mash distances revealed similar trends: the phylogroup D exhibited the highest genetic diversity, followed by F, E, and then by the most abundant groups – A, B1 and B2 – which all have similar levels of diversity (Supplementary Fig. 7). The phylogroup G was the least diverse, but it is also poorly represented in our dataset (33 genomes from three STs). Overall, genetic diversity is proportional to the depth of the phylogroup, i.e. the average tip-to-MRCA distance, except for phylogroup F which is more diverse than expected (Fig. 2b). These results suggest that genetic diversity varies between phylogroups and that within phylogroups it is strongly affected by the time of divergence since the most recent common ancestor.

**Fig. 2:**
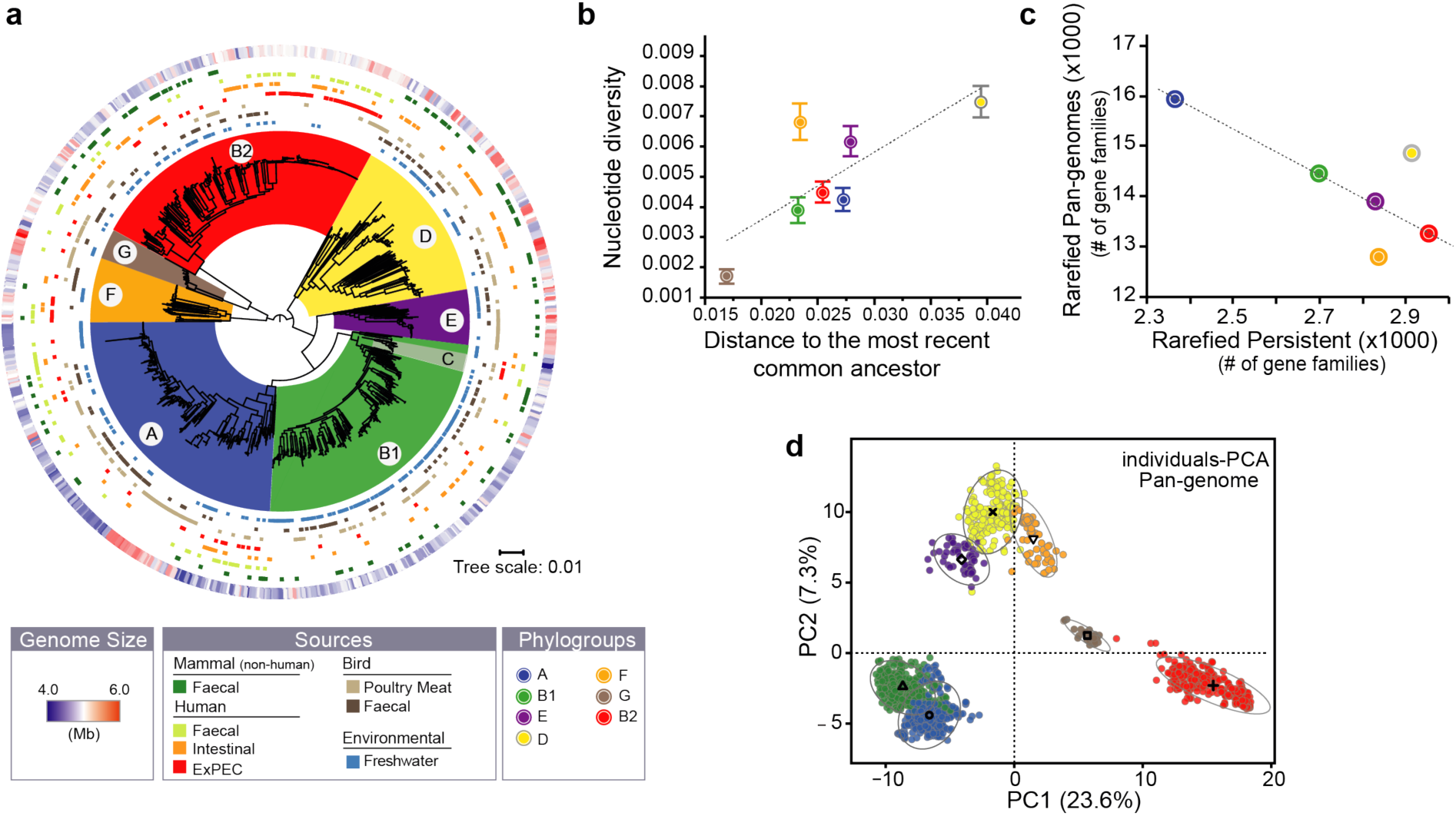
The genetic and ecological structure of Australian *E. coli* population. **(a)** Phylogenetic tree of *E. coli* rooted using the genomes of other *Escherichia* (not shown for clarity). From the inside to the outside: the 7 main phylogroups (arcs covering the tree), the source of each genome (seven rows), and the size of the genomes (outer row, see insert legend). **(b)** Association between the nucleotide diversity per site (Pi, average and s.e) within phylogroup and their distance to their most recent common ancestor (MRCA). In each pylogroup, we averaged the nucleotide diversity (*π)* obtained for 112 core-genes, and the length branches (from tip-to-MRCA) of the species tree. **(c)** Association between the rarefied pan- and persistent-genomes in each phylogroup. We used 1,000 permutations (genomes orderings) of 50 randomly selected genomes (rarefied datasets) to compute the pan- and the persistent-genomes in each phylogroup (ignoring the G group), and then averaged the results. **(d)** Principal component analysis of the pan-genome (matrix of presence/absence of each gene family across genomes). Each dot corresponds to a genome in the two first principal components (PC). The ellipse (90%) and barycenter of each phylogroup are reported. The percentages in the axis labels correspond to the fraction of variation explained by the PC. Panels (b), (c), and (d) have the same color code as (a).

The sets of genomes of each phylogroup have large and open pan-genomes (Supplementary Fig. 8 and Supplementary Table 2). The sizes of these pan-genomes differ widely across phylogroups and are partly correlated to the number of genomes in the phylogroup, explaining why the phylogroup G has the smallest pan-genome (Supplementary Fig. 8). To control for the effect of sample size, we computed pan-genomes from 1,000 random samples of 50 genomes for each phylogroup (ignoring the few strains of the G phylogroup, Fig. 2c and Supplementary Table 2). This revealed larger pan-genomes for phylogroups A, D, and B1 followed by E, B2 and F. Intriguingly, the larger the pan-genome of a phylogroup, the smaller the fraction of its genes that are part of the persistent genome (Fig. 2c). This suggests that differences of pan-genome sizes across phylogroups are caused by different rates of gene turnover in certain phylogroups. They affect all types of genes, even those at high frequency in the species.

To quantify the similarities in gene repertoires, we analyzed the GRR values between phylogroups. The smallest values were observed when comparing B2 strains with the rest (Supplementary Fig. 10). Accordingly, a principal component analysis of the presence/absence matrix of the pan-genome shows a first axis (accounting for 23.6% of the variance) clearly separating the B2 from the other phylogroups (Fig. 2d). This shows that gene repertoires of B2 strains are the most distinct from the other groups, even if B2 is not a basal clade in the species tree. Hence, phylogroups differ in terms of their gene repertoires and in their rates of genetic diversification.

### Mobile genetic elements drive rapid initial turnover of gene repertoires

Different mechanisms can drive the rapid initial diversification of gene repertoires. Mobile genetic elements encoding the mechanisms for transmission between genomes (using virions or conjugation) or within genomes (insertion sequences, integron cassettes) are known to transfer at high rates and be rapidly lost^49–51^. We detected prophages using VirSorter^52^, plasmids using PlaScope^53^, and conjugative systems using ConjScan^54^ (Supplementary Figs. 11-13). These analyses have the caveat that some mobile elements may be split in different contigs, resulting in missed and/or artificially split elements. This is probably more frequent in the case of plasmids, since they tend to have many repeated elements^55^. Only two genomes lacked identifiable prophages and only 9% lacked plasmid contigs. We identified 929 conjugative systems, with some genomes containing up to seven, most often of type MPF_F_, the type present in the F plasmid. On average, prophages accounted for 5% and plasmids for 3% of the genomes (Fig. 3a). Together they account for more than a third of the pan-genomes of each phylogroup. We also searched for elements capable of mobilizing genes within genomes: Insertion Sequences, with ISfinder^56^, and Integrons, with IntegronFinder^57^. Even if ISs are often lost during sequence assembly, some genomes had up to 152 identifiable ISs representing ∼1% of the genome (Fig. 3a and Supplementary Fig. 13). A fourth of the ISs were in plasmids and very few were within prophages. We found integron integrases in 14% of the genomes, usually in a single copy. It is interesting to note that even if the frequency of each type of MGE varies across strains, each of them is strongly correlated with the frequency of the other elements (Fig. 3b). Hence, the typical *E. coli* genome has at least one transposable element, a prophage and a plasmid, the key tools to move genes between and within genomes. When genomes are enriched in one type of MGE, they tend to get simultaneously enriched in the remaining MGEs.

**Fig. 3:**
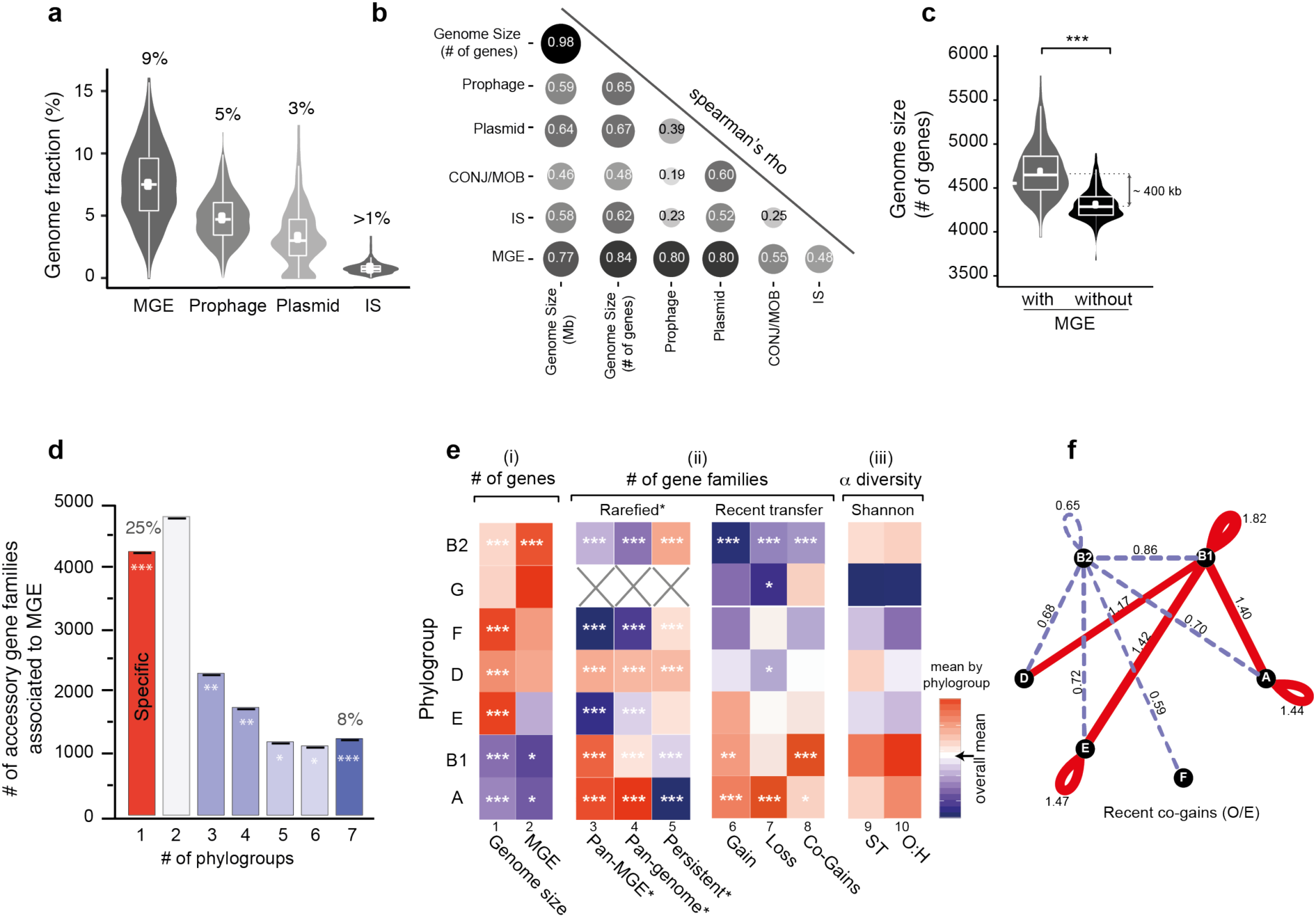
Genetic diversification across phylogroups. **(a)** Percentage of genes associated with MGEs per genome (sum in first graph). **(b)** Spearman’s rank correlation matrix between the number of genes related to MGE (altogether or individually) and the genome size (in Mb and number of genes). Color intensity and the size of the circle are proportional to the correlation coefficients. All values are significantly positive (P<10^-4^). **(c)** Differences in genome size when including or removing gene families associated to MGE (Wilcoxon test, P<10^-4^). **(d)** Number of accessory gene families associated to MGE present in one (i.e., phylogroup-specific) to seven phylogroups. The color code used corresponds to the Z-score obtained for the observed number (O) with respect to the random distribution (E) (see Methods) for each case with a color code ranging from blue (under-representation) to red (over-representation). The level of significance was reported: |Z-score|: * ([1.96-2.58[), ** ([2.58-3.29[, ***([3.29). **(e)** Heatmap where a cell represents the deviation (the difference) of the phylogroup to the rest. All values were standardized by column. The color code ranging from blue (lower) to red (higher), with white (overall mean). The level of significance of each ANOM test was reported: * (P<0.05), ** (P<0.01), *** (P<0.001). **(f)** Network of recent co-occurence of gains (co-gains) of accessory genes within and between phylogroups. Nodes are phylogroups and edges the O/E ratio of the number of pairs of accessory genes (from the same gene family) acquired in the terminal branches of the tree. Only significant O/E values (and edges) are plotted (|Z-score|>1.96). Under-represented values are in dash blue and over-represented in red (see Methods).

What is the effect of these MGEs in the dynamics of *E. coli* genomes? First, the acquisition of MGEs affects the size of the genome. Those identified in this study account for ∼8% of the genome size (Fig. 3c and Supplementary Fig. 14). Accordingly, the number of genes associated with MGEs was strongly correlated with genome size for every type of element (Fig. 3b). Second, MGEs increase the variability of genome sizes, since removing them decreases the coefficient of variation of the size of gene repertoires by 34% (expected increase of 4% under a Poisson model, Fig. 3c). Third, the increase in variance in genome size caused by MGEs is amplified by their short persistence times in the genome. No MGE-associated gene family is sufficiently frequent to be part of the persistent genome, and most (85%) are present in less than 1% of the genomes. For example, 41% of the IS gene families are singletons (Supplementary Fig. 14). Adaptive genes acquired through the action of MGEs may become fixed in populations, but the lack of fixation of recognizable MGEs suggests that the long-term cost of MGEs themselves is significant and/or their contribution to fitness is low (or temporary).

**Fig. 4:**
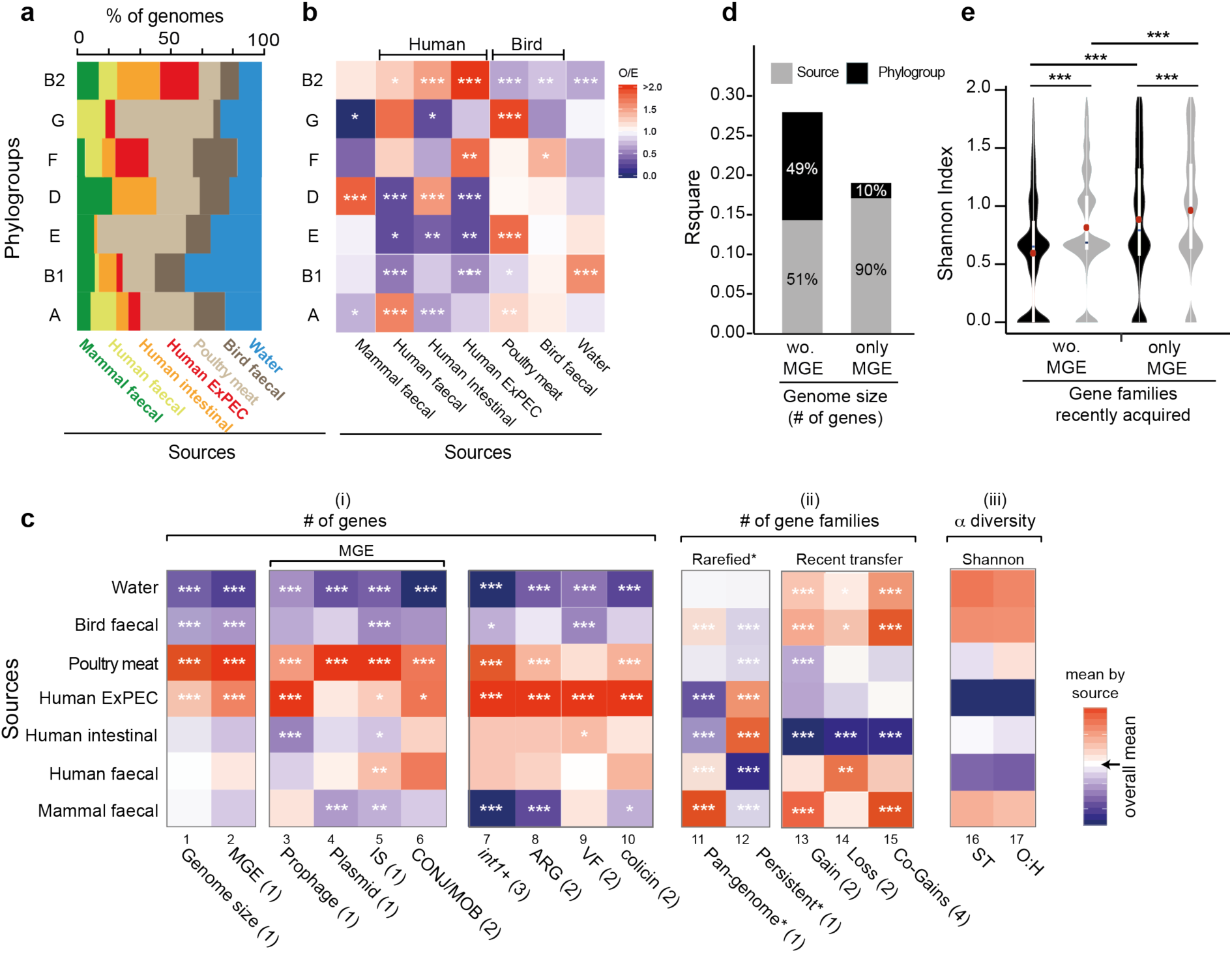
Genetic diversification across sources. **(a)** Distributions of the sources in each phylogroup. **(b)** Association between phylogroups and sources. The ratio of the number of observed (O) genomes divided by the expected (E) number was reported for all comparisons with a color code ranging from blue (under-representation) to red (over-representation) (Fisher’s exact tests performed on each 2*2 contingency table). **(c)** Heatmap showing the associations between isolation sources and a number of traits. Each cell indicates the deviation (the difference) to the overall mean (in white). All values were standardized by column. By default, tests used standard ANOM (1). In presence of deviations from Gaussian distributions, we used non-parametric ANOM tests (2). We used ANOM for proportions (3). We represented the (O/E) ratio of the co-occurrence of gene pairs recently acquired (Co-gains) in each phylogroup with the same color code as in panel (b) (4). **(d)** Contribution of each variable (phylogoup and source) to the variance explained by the stepwise multiple regressions of genome size (for the component of MGEs or the remaining genome) on phylogroup and the isolation source. **(e)** Differences in diversity of gene families recently acquired across phylogroups (in black) and sources (in grey) for gene families associated to MGE or the remaining gene families (Wilcoxon tests, red dots (means)). In all panels : the level of significance of each test was reported: * (P<0.05), ** (P<0.01), *** (P<0.001).

Is the distribution of MGEs associated with phylogroups leading to preferential paths of gene transfer? It has been suggested that homologous recombination is much rarer between than within phylogroups^18^. To test if this applies to the transfer of MGEs, we analyzed the distribution of the pan-genome gene families that are part of MGEs (excluding singletons, for the separate analysis of prophages and plasmids, see Supplementary Fig. 15). Even if these genes are at low frequency in the pan-genome and are observed in a single phylogroup more often than expected by chance (Z-score>20, see Methods), 75% of the phage and plasmid gene families were found in more than one phylogroup and 8% were found in all phylogroups (usually at low frequency, Fig. 3d). Accordingly, the number of gene families present in two to six phylogroups is barely lower, even if significantly so, than expected by chance. These results suggest that there is frequent transfer of MGEs across the different phylogroups. To test this hypothesis more precisely, we used Count to infer gene gain and loss events in the phylogenetic tree of the species (see Methods). We found that half of the recent gene acquisitions, i.e., those that took place at the level of the terminal branches of the species tree, are in families of genes of MGEs. Conversely, the acquisitions at the terminal branches correspond to 40% of the MGE genes of the species. Hence, MGEs are key players in genome diversification at the micro-evolutionary scale. They are transferred across phylogroups and many of them, even if present in several strains, were acquired independently and have just arrived in their host genome.

One might expect more genetic diversity in phylogroups with more MGEs and larger genomes. In apparent agreement with this hypothesis, genomes from phylogroups A and B1 are significantly smaller than the others (Fig. 3e, col 1, ANOM tests, P<10^-3^) and have fewer MGE-associated genes (Fig. 3e, col 2, ANOM tests, P<0.05). However, these phylogroups also have the largest diversity of gene families associated to MGEs (Fig. 3e, col 3, in both the full and rarefied datasets, both ANOM tests, P<10^-3^), *i.e.* they encode fewer but more diverse MGEs. Furthermore, the phylogroups A and B1, in spite of having among the most recent common ancestors of the phylogroups (Fig. 2b), have the largest pan-genomes, the smallest persistent genomes, and the largest diversity of STs, and serotypes (Fig. 3e, in both the full and rarefied datasets, cols 4,5,9,10, ANOM tests, P<10^-3^). This intriguing pattern suggests that the smallest genomes have the highest turnover of genes, not the lower rates of transfer. To test this hypothesis, we took the quantification of gene gains and losses at the terminal branches of the species tree and computed the number of these events per phylogroup. We found that phylogroups A and B1 have the highest number of gene gains and losses per terminal branch (Fig. 3e, cols 6-7). In parallel, we quantified the number of recently acquired (terminal branches) gene pairs (co-gains) from the same gene family within a phylogroup (Fig. 3e, col 8) and between phylogroups (see Methods, Fig. 3f). The results were represented as a graph where the edges represent significantly fewer (dashed lines) or higher (solid lines) number of co-gains than expected by chance. We found that phylogroup B1 has significantly more co-gains of genes with other phylogroups than expected, while the inverse was observed for phylogroup B2. We reached similar results when considering only the co-gains associated with MGEs (Supplementary Fig. 15). These results are consistent with the separation of the B2 phylogroup from the others in the PCA analysis (Fig. 2d). They show that such separation is due to lower rates of transfer in B2, which leads to fewer co-gains within the phylogroup and between this and the other phylogroups. In summary, phylogroups differ in terms of their genome size and in their rates of genetic diversification, the two traits being inversely correlated within the species.

### Not everything is abundant everywhere: the interplay between phylogroups and sources

Frequent horizontal transfer across phylogroups could result in adaptation being independent of the strain genetic background, if there is a lack of epistatic interactions. While we observed that all isolation sources have strains from all phylogroups (Fig. 4a), different phylogroups are typically over-represented depending on the source (Fig. 4b). These observations match previous studies^3^, and suggest strong associations between the phylogenetic structure of populations and the natural habitats of strains.

How much of the variability in gene repertoires is explained by the source of isolation of the strains? Genome sizes vary significantly across isolation sources. Strains isolated from poultry meat had the largest average genomes, followed by ExPEC strains. In contrast, strains from wild birds’ feces and freshwater had the smallest genomes (Fig. 2a and Fig. 4c, col 1, ANOM tests, P<10^-3^). We showed above that genome size also varies across phylogroups. To understand the relative role of the two variables, isolation source and phylogroup, we made two complementary analyses. First, we compared the genome size of strains from different sources within each phylogroup. Even if the statistical power was sometimes low, this revealed trends similar to the ones observed across phylogroups (Supplementary Fig. 17). Second, we used stepwise multiple regressions to assess the effects of phylogroup and the strains’ source on its genome size. Both variables contributed significantly, and in almost equal parts, to the statistical model and together explained 36% of the variance (R^2^=0.36; P<10^-4^, Supplementary Table 3). We found similar results after removing MGE-associated genes (Fig. 4d and Supplementary Table 4). We conclude that both isolation source and phylogroup are equally associated with genome size.

Adaptation to a habitat depends on HGT, which is driven by MGEs. Hence, we studied the distribution of MGEs in relation to isolation sources. There are fewer MGE-associated genes in strains isolated from freshwater and wild birds’ feces, which have smaller genome sizes, and more in strains from ExPEC and poultry meat (Fig. 4c, col 2, ANOM tests, P<10^-3^, and Supplementary Table 5). We observed similar trends within each phylogroup even if the statistical power was low (Supplementary Fig. 17). The analysis of the relative contribution of phylogroups and isolation sources to the number of MGE-associated genes showed that the source of the strain accounted for the vast majority of the explained variance (90%, full model: R^2^=0.19; P<10^-4^, Fig. 4d and Supplementary Table 6). Accordingly, the number of MGE-associated gene families specific to a given source was higher than expected (Z-score >17, Supplementary Fig. 15), and nearly one third of these source-specific families were observed in multiple phylogroups. When we focused on the number of co-occurring recently acquired gene pairs (encoding for MGE or not), we found that they are more frequent within most of the isolation sources than expected by chance (Fig. 4c, col 15, see Methods). These results suggest that the contribution of MGEs to genome size is primarily driven by isolate source rather than phylogroup membership.

The previous result could arise from preferential co-gains of MGEs in an isolation source relative to a phylogroup. To test this hypothesis, we used the results from Count and built a matrix where for each gene family we indicate the acquisition or not of a gene in a terminal branch of the phylogenetic tree. We then compared the clustering of these recent acquisitions by phylogroup and by isolation source using Shannon indexes (see Methods). If the hypothesis is correct, we expected higher clustering (lower diversity) across sources than across phylogroups. We observed slightly higher clustering across phylogroups than across sources, both for MGE-associated and for the other genes (Fig. 4e). We conclude that the contribution of MGEs to genome size depends largely on the isolation source but that this does not reflect systematic co-gains of MGEs in the same source.

It is tempting to speculate that the association between the number of MGE-associated genes and isolation sources reflects selection for the acquisition of locally adaptive functions transferred by these MGEs. To test this, we searched for the presence of antibiotic resistance genes (ARGs) in our dataset using the reference databases. Many of these ARGs were in integrons (∼3 per integron), which is well documented^58^, and genomes carrying integrons had more ARGs than the others (Wilcoxon test, P<10^-4^, Supplementary Fig. 18). Expectedly, integrons and ARGs were more prevalent in ExPEC and in poultry meat isolates (Fig. 4c, cols 7-8) and Supplementary Table 5). Similar results were observed in the analyses at the level of each phylogroup (Supplementary Fig. 18). The clear association of integrons and ARGs with human (or domesticated animals) isolates of *E. coli* independently of the phylogroups’ genetic background reinforces the idea that source-specific MGEs provide locally adaptive traits.

To complement the previous results, we searched for the presence of other factors known to be adapative under specific conditions: virulence factors involved in antagonistic interactions with humans and colicins involved in intra-specific competition. Virulence factors (VFs) from VFDB are more prevalent in human strains with an excess in ExPEC isolates (ANOM test, P<10^-3^) and less frequent in strains isolated from freshwater and wild birds’ feces (ANOM test, P<10^-3^, Fig. 4c, col 9). While VFs are more concentrated in phylogroups B2, D, E and F (ANOM test, P<10^-2^) as previously shown^37^, the trends regarding isolation sources are conserved within each phylogroup (Supplementary Fig. 19). In particular, within phylogroup B2, only human strains have a significantly higher average number of VFs (Supplementary Fig. 19) reinforcing previous results^26^. We also analyzed colicin gene clusters, which are agents of bacterial antagonistic competition and are often encoded in plasmids^59^. The average number of colicins identified using BAGEL3^60^ (some of which are also included in VFDB) depends on the phylogroup of the strain, from an average of 2.8 genes in B2 strains to 0.4 in B1 strains. Interestingly, the water isolates have the fewest colicin genes, presumably because free diffusion of these proteins in water makes them inefficient tools of bacterial competition (Fig. 4c, col 10 and Supplementary Fig. 19). Thus, local adaptations resulting from the acquisition of novel genes by HGT, involving antagonistic interactions with other bacteria or with the host, are associated preferably with certain phylogroups. This may result from specific genetic interactions in the different genetic backgrounds.

*E. coli* has usually been regarded as a contaminant from animal, mostly human, sources and used to test water quality. Yet, recent data suggests that some strains could inhabit aquatic environments^61^. Given the contrast between the primary and secondary habitats of *E. coli*, respectively guts of endotherms and aquatic environments, this would imply marked differences between the 285 freshwater strains and the others. Indeed, our results show that these strains are systematically different. They are over-represented in phylogroup B1 (43%), a phylogroup under-represented in all other sources of isolation (Figs. 2a,4b). On the other hand, they are under-represented in B2 (13%), a phylogroup over-represented in strains isolated from humans (this study) and other mammals^2^. The genome size of freshwater strains’ is the smallest among all groups of isolates and across phylogroups (Fig. 4c, col 1, Supplementary Fig. 17). Importantly, these strains show average pan-genome sizes in the rarefied dataset, suggesting that adaptation is not exclusively due to genome reduction (Fig. 4c, col 11). This is also supported by the high number of gains and losses observed (Fig. 4c, cols 13,14), although these genomes have the fewest MGEs and often lack plasmids (Fig. 4c, cols 2-6). Consistent with adaptation to this habitat, they have the smallest number of antibiotic resistance genes, virulence factors, and bacteriocins (Fig. 4d, cols 7-10) and Supplementary Fig. 18,19). In contrast, these strains show the highest diversity of STs and O:H serotypes (Fig. 4c, cols 16,17, and Supplementary Table 5). The extreme genomic traits of isolates from water strongly suggest they are not the result of recent fecal contamination from other sources. Instead, they strongly suggest that these strains have changed to adapt to water environments. If so, this seems to have involved extensive horizontal gene transfer concomitant with streamlining, i.e. a high turnover of gene repertoires that resulted in genomes smaller than the average.

## Discussion

Many of the recent advances in the understanding of *E. coli* evolution focused on clinical isolates and placed a lot of emphasis on virulence and antibiotic resistance in a few clinically important lineages^62–67^. Yet, most strains of the species are commensal. Hence, most of the evolution of the species takes place in biotic contexts not associated with pathogenesis. Furthermore, while a lot of attention has been placed on the rates of homologous recombination in the chromosome of the species, it is now clear that HGT drives the evolution of virulence^12, 42, 68, 69^ and antibiotic resistance^70–72^ in pathogenic strains as well as that of many other traits in commensal strains^12^. For example, MGEs were recently shown to be more important than point mutations for the colonization of the mouse gut by *E. coli* commensals^73^. Here, we aimed at providing a global picture of the evolution of the *E. coli* genomes with an emphasis on the variation of gene repertoires in strains from a variety of sources (environmental and geographic) across a single continent. This allowed us to study the joint effect of population structure and habitat on the variation of gene repertoires. Our study focused on *E. coli* isolates from Australia, but its genetic diversity was higher or comparable to other worldwide genome datasets, and its population structure was consistent with previous works^16, 40, 74^. This indicates that what we have observed is likely to be representative of the species as a whole. It also confirms previous reports of the large genetic diversity of the species and of the planetary circulation of all major lineages^39, 45, 75^. Finally, the functional annotation of the pan-genome shows that in spite of over 375,000 papers citing *E. coli* in PubMed in 2019, we are still far from having discovered the full genetic diversity of *E. coli* and from knowing the function of many of its most frequent gene families.

We started our study by quantifying gene repertoire diversification, which we found to follow a two-step dynamics. The very rapid initial diversification, where GRR quickly decreases to ∼90%, implicates substantial heterogeneity in terms of gene repertoires for strains that are from the same sequence type and are almost identical in the sequence of persistent genes. Some of this divergence may be due to genome sequencing or assembling artifacts producing singletons and thus inflating pan-genomes. Yet, we have annotated all genomes in the same way. We also confirmed key results by excluding singletons, and showed that singletons represent only ∼0.5% of a typical genome and that many of them have homologs in the databases. The frequency of singletons is only weakly correlated with the number of contigs in draft assemblies, a further sign that they are not just caused by sequencing or assembly issues (Supplementary Notes). Furthermore, our analysis of ancestral genomes showed that a large fraction of well-known MGEs, including phages, ISs and plasmids, were acquired very recently (inferred acquisition at the terminal branches of the phylogenetic tree). Some of these are singletons, whereas others are present across many phylogroups. They contribute directly to the rapid divergence of gene repertoires between separating lineages. Previous population genetics models applied to other clades observed the existence of genes that have rapid turnovers in genes^76, 77^. Our results show that frequent acquisition of MGEs drives rapid diversification of gene repertoires even between strains that are almost indistinguishable by classical typing schemes.

Following the abrupt initial loss of GRR between diverging lineages, we observed that the similarity of gene repertoires decreases linearly with time. Hence, it does not follow the negative exponential distribution that we proposed a decade ago^42^, which was based on a very small set of genomes that precluded the identification of the change of dynamics at small patristic distance. This change of dynamics resembles the accumulation of non-synonymous mutations in genes under weak purifying selection. Comparisons between closely related strains reflect almost neutral accumulation of recent events whereas differences between distant strains are driven by purifying selection with occasional fixation of adaptive events^78, 79^. In the present context, this suggests that either many integrations of genetic material are slightly deleterious or that there is rapid deletion of neutral genes. The first hypothesis is consistent with the fitness costs associated with the acquisition of many MGEs^80–82^, and with our observation that most MGEs present in a genome were very recently acquired. The second hypothesis is consistent with the previous works suggesting the existence of mechanistic biases towards gene deletion in bacteria ^83, 84^. Once most the recent transfer has been purged, by natural selection or gene deletion biases, GRR decreases linearly with divergence time and shows large variance around the regression line. The large variance indicates that some distantly related bacteria may have more similar gene repertoires than bacteria within the same sequence type. Importantly, the analysis does not suggest the existence of a point beyond which relatedness and gene flow change abruptly. Hence, these results do not suggest incipient sexual isolation within the species from the point of view of horizontal gene transfer. The analysis of gene flow associated with B2 strains should be placed in this context, it shows that this particular phylogroup has many MGEs and large genomes, but is recently exchanging less genetic material with strains from its own and from other phylogroups. This has placed it apart from the other phylogroups in terms of gene repertoires and in terms of preferential habitats.

The rapid evolution of gene repertoires by HGT is consistent with the observation that plasmids, prophages and ISs are almost ubiquitous among *E. coli*. These elements contribute to the genome size and especially to its variability across strains, which supports our previous results^50, 85^. While most MGEs are quickly lost from lineages, or drive the lineage extinct, the large influx of such elements can bring adaptive accessory traits such as antibiotic resistance genes^71^ and virulence factors^86, 87^. They also pave the way for cooption processes^88^. The contribution of the MGE genes to genome size across the species is more strongly associated with the isolation source of the strains than with the phylogroup. However, the recent co-acquisition of MGEs by different strains is also associated with the phylogroup. This is consistent with a scenario where the abundance of MGEs in a genome is strongly dependent on the habitat, but their diversity also depends on the phylogroup. Since most MGE genes arrived in the genome very recently, this suggests that habitat exerts a strong constrain on the flow of gene exchanges across *E. coli* strains, in line with the view that bacteria exchange more genes with those they coinhabit^89, 90^.

The need of favorable genetic backgrounds for certain local adaptation processes could explain the observed over-representation of some phylogroups in certain isolation sources. Virulence factors and antibiotic resistance genes provide relevant examples. In our dataset, the plasmids encoding virulence factors are often conjugative and should be able to circulate widely, but the virulent clones often concentrate in only a few phylogroups. Selection for antibiotic resistance is expected to be higher in the virulent clones, because these are the most targeted in the clinic. Hence, they endure stronger selection to keep the ARGs arriving in MGEs. These causal links result in preferential associations of genetic backgrounds with virulence factors and ARGs, and therefore with the frequency of pathogens in a given phylogroup. How much of these trends are due to epistatic interactions between novel genes and the genetic background and how much is due to availability of specific genes by horizontal transfer in certain sources remains to be quantified. In conclusion, these results contribute to explain why epidemiological clones tend to emerge from specific phylogenetic groups even in the presence of massive horizontal gene transfer.

Genetic diversity, created by HGT, recombination, or mutation, affects a species’ ability to adapt to novel ecological opportunities. The higher the diversity of gene repertoires in a population, the more likely that one of those genes will prove helpful in the face of environmental challenges such as antibiotics. We observed that the generalist phylogroups, such as A and B1, have broader pan-genomes than specialist phylogroups like B2. This was not expected based on their smaller genome sizes or the lower frequency of MGEs in their genomes. We propose that this reflects the high variability of the environments where they circulate and the consequent diversity of local adaptation processes. Phylogroup B2 strains, by comparison, have developed very specific traits that may let them take advantage of some particular resources, e.g. they are better adapted to mammal gut environment^2^. This has resulted in large genomes that have diverged more from the other *E. coli*, as revealed by the PCA analysis, but that are overall more conserved (largest persistent-genome, smaller pan-genomes, fewer recent gene acquisitions). Altogether, these results suggest that the habitat and the phylogenetic structure jointly determine the size of genomes. The results also suggest the hypothesis that the large genomes of some phylogroups, like B2, are caused by a relative decrease in the rate of gene loss, not by an increase in the rate of gene gain.

The integration of information on gene repertoires and population structure in strains sampled from diverse sources can shed light on the origin of environmental strains. This is illustrated by the identification of genomic traits in freshwater *E. coli* isolates that are very different from the average traits of the species and that suggest adaptation of certain lineages to this environment. For bacteria, freshwater environments are much more nutrient poor than the guts of endotherms, and it’s interesting to note that strains associated with this environment have more streamlined genomes. This may represent at the micro-evolutionary scale, an adaptation similar to that observed in other bacteria adapted to poor nutrient environments that have small genomes and few MGEs^91, 92^. These results are also consistent with recent studies showing that *E. coli* B1 strains can persist longer in water than strains of the other phylogroups, and that B1 persistent strains in water often encode very few virulence factors and antibiotic resistance genes^7, 33, 34^. Interestingly these strains have been shown to be able to grow at low temperatures^7^. The prevalence of B1 isolates has been observed in other environmental samples, such as drinking water or plants^93^. The characteristics observed in freshwater isolates might be general to this environment, since they were observed in strains from the B1 and from other phylogroups (Supplementary Figs. 16-18). If some *E. coli* lineages are indeed adapted to freshwater this radically changes the range of environments from where they can acquire novel genes and the selection pressures that shape their subsequent fate. This finding also implies that environmental isolates are not necessarily the result of source-sink dynamics where *E. coli* strains evolve in relation to selection pressures linked to the host and environmental strains are just sinks where such strains find evolutionary dead-ends. Instead, the environment outside the host could have a significant impact on the evolution of *E. coli* subsequently colonizing human hosts.

## Materials and Methods

### Strains

We used different collections of *E. coli* strains recovered in Australia between 1993 and 2015 (for a more detailed description, see Supplementary Note and Supplementary Dataset1). The subset of strains selected for whole genome sequencing includes : (1) *faecal strains* isolated from various birds (N=195 strains), non-human mammals (N=135), and humans living in Australia (N=93); (2) *clinical strains* isolated during intestinal biopsies of patients with inflammatory bowel disease (N=172), or corresponding to human ExPEC strains collected from urine or blood (N=112); (3) *poultry meat strains* isolated from chicken meat products from diverse supermarket chains and independent butcheries (N=283); (4) and *freshwater strains* isolated from diverse locations across Australia (N=285).

### Sequencing

Of the 1,304 isolates, 70 were sequenced at Broad institute using the Roche 454 GS FLX system, 70 were sequenced by GenoScreen (Lille, France) using the HiSeq2000 platform and the rest were sequenced at the Australian Cancer Research Foundation (ACRF) Biomolecular Resource Facility (BRF) of the Australian National University using the Illumina MiSeq platform.

### Assembling

Paired-end read files were processed and assembled with CLC Genomics Workbench v.9.5.3 (Illumina) using their *de novo* assembly algorithm with default parameters. All genomes sequenced by the Broad institute were available into the NCBI Assembly (www.ncbi.nlm.nih.gov/assembly/) or SRA (www.ncbi.nlm.nih.gov/sra/) databases. While, the rest of the assemblies was deposited into the European Nucleotide Archive (PRJEB34791). The accession number of each genome is reported in Supplementary Dataset1.

### Datasets

We used 4 datasets in this study. (1) The ***Australian dataset*** described above is the main dataset. (2) ***RefSeq dataset***: We retrieved 370 *E. coli* complete genomes from GenBank Refseq (available in February 2018). (3) ***ECOR dataset***: We retrieved 72 draft genomes of the *E. coli* reference (ECOR) collection from DDBJ/ENA/GenBank^48^. Strains in this collection were isolated from diverse hosts and geographic locations and have been used for more than 30 years to represent the phylogenetic diversity of *E. coli* as they have been selected from over 2,600 natural isolates based on MLEE data^17^. (4) ***Outgroup dataset***: We retrieved 65 other closely related *Escherichia* genomes from ENA/GenBank and sequenced 21 others on the Illumina MiSeq platorm (assembled as described above). They belong to Clade I (N=14), Clade II (N=2), Clade III (N=8), Clade IV (N=2), Clade V (N=14), *E. fergusonii* (N=8) and *E. albertii* (N=38) species. Only five of them were complete, others were draft genomes. In this study, these genomes (called hereafter *outgroup* genomes) were only used to root the Australian *E. coli* species tree. The general genomic features and the sequencing status of these 1,832 genomes are reported in Supplementary Dataset1.

### Data formatting

In an attempt to overcome the bias from different annotations all genomes of the four datasets were annotated using Prokka v.1.11^94^ which provided consistency across the entire datasets (with hmmer v.3.1b1, aragorn v.1.2.36, barrnap v.0.4.2, minced v.0.1.6, blast+ v.2.2.28, prodigal v.2.60, infernal v.1.1, ncbi_toolbox v.20151127, and signalp v.4.0). We performed three quality controls on genomic sequences of Australian and outgroup datasets (see Supplementary Note). A total of 10 *E. coli* draft genomes and one genome from clade V failed at least one of these tests and were removed from further analysis, leading to a final dataset of 1,294 Australian *E. coli* genomes and 87 outgroup genomes. The main characteristics of each draft genome are reported in Supplementary Dataset1.

### E. coli typing

***Phylogroup***. The phylogroup of each *E. coli* genome (from ECOR, RefSeq, and Australian datasets) was determined using the *in silico* ClermonTyping method^20^. ***Multilocus sequence typing*** (***MLST).*** Sequence type (ST) was identified by the MLST scheme of Achtman^10^ using mlst v.2.16.1 (https://github.com/tseemann/mlst). We assigned STs for a large majority of genomes, i.e., for 99%, 96% and 97% of the ECOR, RefSeq and Australian genomes resp. ***Serotype***. Serotype (O- and H-genotypes) was inferred with the EcOH database^95^ using ABRicate v.0.8.10 (https://github.com/tseemann/abricate). Currently there are 220 *E. coli* O-groups and 53 H-types described in this database. While 99% of Australian genomes had H-group assigned, only 57% had O-group assigned even if *wzm*/*wzt* and *wzx*/*wzy* genes are present. All these results are reported in Supplementary Dataset1.

### Nucleotide diversity

The ***nucleotide diversity*** of the three datasets, *i.e.*, ECOR, RefSeq and Australian, was computed from the multiple alignments of 112 core gene families present in all *E. coli* genomes of these three datasets, (see below), using the diversity.stats function from the *PopGenome* v.2.6.1 R package^96^. We also used these 112 core gene families to assess the nucleotide diversity for each phylogroup of the Australian dataset.

### ST and O:H diversity

The ***Shannon index*** was computed to assess the diversity of ST and O:H serotypes within each phylogroup and source. For this, we calculated their relative frequency in each group and then applied the function skbio.diversity.alpha_diversity from the *skbio.diversity* v.0.4.1 python package (http://scikit-bio.org/docs/0.4.1/diversity.html).

### Mash distances (M)

***Genome similarity.*** Due to the high cost of computing ANI^97^ via whole-genome alignment, we estimated genome similarity calculating the pairwise Mash distance (M) between all Australian genomes using Mash v.2.0^98^. Importantly, the correlation between the Mash distances (M) and ANI in the range of 90-100% has been shown to be very strong, with M ≈ 1-(ANI/100)^98^. All the resulting Mash distances between *E. coli* genomes are well below 0.05, in agreement with the assumption that they all belong to the same species. The median is 0.027 and the maximal value is 0.04 (Supplementary Fig. 3). **Australian *E. coli* reference genomes**. The Mash distance was strongly correlated to the patristic distance in our dataset (spearman’s rho=0.92, P<10^-4^). We used it to select 100 Australian *E. coli* strains representative of the species’ diversity (called hereafter *reference* genomes). Such *reference* genomes were used to root the Australian *E. coli* tree (to drastically reduce the computational time required to build the rooted tree). To select representative genomes, we performed a hierarchical WPGMA clustering from the Mash distance matrix computed with all Australian *E. coli* genomes, and then we cut it off to have only 100 clusters. In each of these clusters, the genome with the smallest L90 was selected. This *reference* dataset contained all the phylogroups and was composed of: 15-A, 10-B1, 13-E, 39-D, 11-F, 10-B2 and 2-G genomes.

### Identification of pan-genomes

Pan-genomes are the full complement of genes in the species (or dataset, or phylogroup) and were built by clustering homologous proteins into families. We determined the lists of putative homologs between pairs of genomes with MMseqs2 v.3.0^99^ by keeping only hits with at least 80% identity and an alignment covering at least 80% of both proteins. Homologs proteins were then clustered by single-linkage^100^. We computed independently the pan-genome of each dataset, *i.e.*, ECOR, RefSeq, Australian and of the 87 outgroups with the 100 Australian *E. coli* reference genomes. Each pan-genome was then used to compute a matrix of presence-absence of gene families. Hence, gene copy number variations were not taken into account in this part of the study. The alpha exponent of Heap’s Law was used to infer whether a pan-genome is open or closed^46^. Thus, if α (alpha) < = 1, the pan-genome is open. In contrast, α (alpha) > 1 represents a closed pan-genome. This coefficient was computed using the *heaps* function of the *micropan* v.1.2 R package^101^ with n.perm = 1000. Principal component decomposition of the Australian pan-genome, *i.e*, the matrix of presence-absence of protein families was computed using the *prcomp* function from the *stats* v.3.5.0 R package.

The pan-genome of each phylogroup and source was taken from the pan-genome of the species. The pan-genome of the MGE (called **Pan-MGE**) was also taken from the species pan-genome and contained only genes encoding for MGEs.

### Rarefaction of pan-genomes

The number of singletons was strongly correlated to the number of genomes analyzed in each phylogroup (Pearson’s correlation = 0.97, P<10^-4^), indicating that the pan-genomes size depend on the number of genomes analyzed. Thus, to compare genetic diversity across datasets (e.g. phylogroups), we rarefied the genome datasets, *i.e.*, each pan-genome was constructed with the same number of genomes in each comparison. To do this, 1,000 subsets of X genomes (X depending on the analysis, specified in the results section) were randomly selected for comparison in each group, resulting to datasets called hereafter *rarefied* datasets (Supplementary Fig. 8).

### Identification of persistent-genomes

Gene families that are persistent were taken from the analysis of pan-genomes. A gene family was considered as persistent when it was present in a single copy in at least 99% of the genomes. We found 2,486 persistent gene families when considering the 1,294 Australian genomes, representing 52% of the average genome.

### Identification of core-genome

The core genome was taken from the analysis of the pan-genome. A gene family was considered as core if it is present in one single copy in all the genomes. To assess the nucleotide diversity, we built a core-genome with all the genomes of the ECOR, RefSeq, and Australian datasets. It was composed of 112 core gene families. Each gene family was aligned with mafft v.7.222 (using FFT-NS-2 method)^102^, and used to compute the average nucleotide diversity (*π*) in each dataset and within each phylogroup (see above).

### Functional assignment of the pan-genome

Gene functional assignment was performed by searching for protein similarity with hmmsearch from HMMer suite v.3.1b2^103, 104^ on the bactNOG subset of the EggNOG v.4.5.1 database^47^. We have kept hits with an e-value lower than 10^-5^, a minimum alignment coverage of 50% of the protein profile, and when the majority (>50%) of non-supervised orthologous groups (NOGs) attributed to a given gene family pertained to the same functional group (category). The gene families that cannot be classified into any existing EggNOG clusters were grouped into the “unknown” category. Hits corresponding to poorly characterized or unknown functional EggNOG clusters were grouped into the “poorly characterized” category.

### Phylogenetic analyses

We built a rooted phylogeny of the species in two steps. **The phylogenetic species tree of Australian *E. coli*** was reconstructed from the concatenated alignments of the 2,486 persistent proteins of the 1,294 Australian *E. coli* strains. Each of these protein families was aligned with mafft v.7.222 (using FFT-NS-2 method)^102^. At this evolutionary distance the DNA sequences provide more phylogenetic signal than protein sequences. Hence, we back-translated the alignments to DNA, as is standard usage. We built phylogenies from persistent genomes to avoid the loss of signal associated with the small core genomes. When a genome lacked a member of a persistent gene family, or when it had more than one member, we added a stretch of gaps (‘-‘) of same length as the other genes for it in the multiple back-translated alignments. Adding a few “-” has little impact on phylogeny reconstruction^105^. We have not removed recombination tracts from the multiple alignment because this has been shown to amplify errors in determining phylogenetic distances and it usually does not affect the topology of the tree^106, 107^. If determination of the recombination was accurate in our >1,300 genomes dataset, this would have led to the exclusion of almost all the genes. The length of the resulting alignment for the species was 2,298,168 bp. Each tree was computed with IQ-TREE multicore v.1.6.7^108^ under the GTR+F+I+G4 model. This model gave the lowest Bayesian Information Criterion (BIC) among all models available (option –m TEST in IQ-TREE). We made 1,000 ultra-fast bootstraps to evaluate node support (options –bb 1000 –wbtl in IQ-TREE) and to assess the robustness of the topology of each tree^109^.

**The phylogenetic tree of *Escherichia* genus** was inferred from the persistent-genome obtained with the 87 outgroup genomes and the 100 *E. coli* reference genomes (see above) using the same procedure as the species tree. In this case, the persistent-genome is composed of 1,589 proteins families, and the resulting alignment of 1,469,523 bp. The genus phylogenetic tree was extremely well supported: all nodes had bootstrap support higher than 95%. Its topology was consistent with a previous study^110^ (Supplementary Fig. 3c). Then, we used it to precisely root the species tree (Supplementary Fig. 3d).

#### The most recent common ancestor of each phylogroup

We identified the node corresponding to the most recent common ancestor (MRCA) for each phylogroup from the rooted species tree using the *findMRCA* function from the *phytools* v.0.6.44 R package. Then, the subtree of each phylogroup was extracted using the *extract.clade* from the *ape* v.5.2 R package^111^. The distance to the MRCA was computed from the length of branches in each subtree. It corresponds to the average depth (distance from the MRCA) of all genomes (tips) within a phylogroup, and was inferred using the *depthTips* from the *phylobase* v.0.8.6 R package (https://github.com/fmichonneau/phylobase).

### Evolutionary Distances

For each pair of genomes, we computed a number of measures of similarity : 1) The **Patristic distance** was computed from the length of branches in the *Australian E. coli* species phylogenetic tree. The patristic distance is simply the sum of the lengths of the branches that link two genomes (tips) in the tree, and was inferred using the *cophenetic* function from the ape v.5.2 R package^111^. They were computed between all pairs of genomes, of the same ST (*intra-ST*), of different ST (inter-ST) within identical phylogroup, or of different phylogroups (*Inter-phylogroup*). As expected, we found that the *intra-phylogroup (both intra-ST and inter-ST)* patristic distances were significantly shorter than the *inter-phylogroup* (Wilcoxon test, P<10^-4^). 2) **The Gene Repertoire Relatedness index** (GRR) between two genomes was defined as the number of common gene families (the intersection) divided by the number of genes in the smallest genome^112^. It is close to 100% if the gene repertoires are very similar (or one is a subset of the other) and lower otherwise. 3) **The Manhattan index** between two genomes is the number of different gene families. If two genomes have identical gene content, the corresponding Manhattan index is 0. 4) **The Jaccard index** between two genomes was defined as the number of common gene families (the intersection) divided by the number of gene families in both (the union). The Jaccard index between two genomes describes their degree of overlap with respect to gene family content. If the Jaccard distance is 1, the two genomes contain identical protein families. If it is 0 the two genomes are non-overlapping.

To characterize the genetic diversification of each phylogroup of the Australian dataset, we computed the three different standard indexes: the GRR, the Jaccard, and the Manhattan indexes. All these indexes were highly correlated (Supplementary Fig. 9). Thus, only analyses with GRR were reported and illustrated in the main text. Note that we always used the matrix of presence/absence of gene families to compute all these indexes, meaning that multiple occurrences were not considered. This downplays the impact of IS on pan-genome size and makes more conservative estimates of GRR divergence.

### Reconstruction of the evolution of gene repertoires

We assessed the evolutionary dynamics of gene repertoires of the Australian genomes using Count (downloaded in January 2018)^113^ with the Wagner parsimony method. Due to the size of our dataset it was not possible to do the analysis using birth-death models, but our previous analyses revealed very few differences between the two methods in smaller datasets^114^. Wagner parsimony penalizes the loss and gain of individual family members (with relative penalty of gain with respect to loss of 1, option g = 1), and infers the history with the minimum penalty. Thus, from the pan-genome, *i.e.*, the matrix of presence-absence of gene families, and the rooted species tree, Count inferred the most parsimonious gain/loss scenario of each gene family along the tree. At each tree node, Count detailed information about individual families: presence/absence, and family events on the edge leading to the node. Hence, we have reconstructed the gene content of ancestral genome at each node. At each terminal branch, the expected total number of recent acquisitions (HGT) was computed by summing all family-specific gene gains obtained from the edge leading to the tip. Among them, we identified MGE associated genes that were recently acquired in each genome. We applied a similar strategy to identify recent losses.

### Distribution of accessory families across phylogroups (or sources)

We counted the number of MGE-associated gene families across phylogroups (Fig. 3d) or sources (Supplementary Fig. 15). We excluded the singletons from this analysis to avoid over-estimation of the number of families specific to one category. To test if some categories over-represented or under-represented these genes, we made 1,000 simulations. In each simulation, we shuffled the phylogroup (or source) assignment of the genomes while keeping the same number of taxa in each category (phylogroups or sources). Thus, the presence of a gene family in a genome and its frequency in the pan-genome remains the same, only the phylogroup (or the source) of genomes changes. The Z-score obtained for the observed number in the real data with respect to the random distribution (from 1,000 simulations) was reported for each case with a color code ranging from blue (under-representation, Z-score<-1.96) to red (over-representation, Z-score>1.96).

### Recent co-occurrence of gains (co-gains) of gene families within phylogroups

We counted the number of recently acquired gene pairs (co-gains) from the same pan-genome gene family (see above) within and between phylogroups. Recently acquired genes were defined as those inferred as acquired in terminal branches using Count. To test if some phylogroups over-represented or under-represented these co-gains, we compared the observed number (O) within each phylogroup to the expectation (E) given by 1,000 simulations. In each simulation, we shuffle the phylogroup assignment of the taxa (same approach as for the accessory gene families) and count the number of co-gains within and between phylogroups. For each phylogroup, we then divided the number observed in the real data (O) by the average number observed in the simulations (E), and computed the Z-score of the observed number (O) with respect to the random distribution (E). We considered an over(under)-representation significant when Z-score>1.96 (Z-score<-1.96). Note that the O and E numbers had to be previously normalized (divided by the total number of gene pairs, i.e. the sum of pairs within and between phylogroups, in the real data, and in each simulation, resp.). We applied the same approach (i) considering only gene pairs encoding for MGEs (similar result as in Fig. 3), (ii) for sources (instead of phylogroups, Fig. 4).

### Network of co-occurrence of gains (co-gains) of gene families across phylogroups

All co-gains (see above) were split into all possible combinations of phylogroup pairs (21 combinations). To test if these co-gains are over- or under-represented between phylogroups, we compared the observed number (O) between each phylogroup to the expectation (E) given by 1,000 simulations with the same strategy as above. As before, we normalized the observed and expected numbers by the total number of co-gains in each simulation, calculated the (O/E) ratio, and the Z-score of each observed value in the real data with respect to the random distribution (E). The network was drawn using the *igraph* v.1.2.2 R package (https://igraph.org/r/) with the circle layout option, where nodes are phylogroups, edges are (O/E) values for which the Z-score is significantly different from zero. The width of the edges is proportional to the (O/E) value and the color is blue for under- and red for over-representation (Fig. 3f). We applied the same approach considering only gene pairs encoding for MGEs (Supplementary Fig. 16).

### Gene family diversity

We computed Shannon indexes to assess the diversity of each gene family recently acquired (terminal branches) across phylogroups and across sources (Fig. 4e). If diversity is low, this means that acquisitions are clustered by phylogroup or source (depending on the analysis). For this, we calculated the relative frequency of each gene family recently acquired within each phylogroup (vs. each source). It is simply the number of genomes (within a phylogroup) with at least one acquisition divided by the total number of genomes in the phylogroup. We therefore obtained 2 vectors per gene family (one for phylogroups and one for sources) each containing 7 frequencies (for each phylogroup or each source) and then applied for each vector the function *diversity* from the *vegan* v.2.4.6 R package (https://github.com/vegandevs/vegan). If the index is 0, recent acquisitions of genes of the family are limited to a single group (phylogroup or source). The higher the index, the more scattered the acquisitions of the family’s genes are (across phylogroups or sources).

### Statistics

All basic statistics were performed using R v 3.5.0, or *JMP*-13. (i) **Analysis of means**: We used **ANOM** to compare group means to the overall mean, when the data were approximately normally distributed. In cases where the data were clearly non-Gaussian and could not be transformed, we used the nonparametric version of the ANOM analysis, i.e., **ANOM with Transformed Ranks**. It compares each group’s mean transformed rank to the overall mean transformed rank. In both, we used the methods implemented in *JMP*-13. (ii) **Pairwise Wilcoxon Rank Sum Tests** were computed using the *pairwise.wilcox.test* function from the *stats* v.3.5.0 R package. We used the Bonferroni correction during multiple comparison testing. (iii) **Fisher’s exact tests** were computed using the *fisher.test* function from *stats* v.3.5.0 R package. They were performed for testing the null of independence of rows (phylogroups) and columns (sources) in a 2×2 contingency table. (iv) **Correlation coefficients.** Pearson’s and Spearman’s rank correlation rho were computed using the *cor* function from *stats* v.3.5.0 R package. The correlation matrices were represented using the *corrplot* v.0.84 R package (https://cran.r-project.org/web/packages/corrplot/index.html). (v) **Smooth regression**: We used the generalized additive model (*gam)* smoothing method from the *mgcv* v.1.8.23 R package (https://cran.r-project.org/web/packages/mgcv/index.html). (vi) **Stepwise multiple regressions** were computed with *JMP*-13. This standard statistical method consists in a stepwise integration of the different variables in the regression by decreasing order of contribution to the explanation of the variance of the data^115^. We used the forward algorithm and the BIC criterion for model choice in the multiple stepwise regressions. The P-values associated with each variable were assessed using an F-test.

### Identification of Mobile Genetic Elements (MGEs)

**Prophages:** Prophages were predicted using VirSorter v.1.0.3^52^ with the RefSeqABVir database in all genomes from Australian and RefSeq datasets, as a control. The least confident predictions, *i.e.*, categories 3 and 6, were excluded from the analyses in both datasets. The prophage-associated regions in drafts are more numerous and shorter than in complete genomes (Supplementary Fig. 11). These results reveal that such regions are sometimes split in assemblies. In complete genomes, the cumulative size of the prophage-associated regions (X) is highly correlated with the number of prophages (Y) present in the genomes (Y=1.2923362 + 1.6767.10^-5^ X, R^2^=0.91, P<10^-4^, Supplementary Fig. 11). Hence, we used this linear equation to estimate the number of prophages in drafts using the cumulated size of prophage regions in the draft genomes. **Plasmids**: In the RefSeq dataset, all the extrachromosomal replicons were considered as plasmids. In the Australian dataset, plasmid sequences were identified using PlaScope v.1.3^53^ with the database dedicated to *E. coli*. PlaScope provides a method for plasmid and chromosome classification of *E. coli* contigs. It has the specificity to select a unique assignment to each contig of a draft genome to plasmid, chromosome or unclassified. The number (∼16, max: 124) and size (∼9 kb, max: 166 kb) of contigs predicted as plasmid were highly variable (Supplementary Fig. 12) in the Australian dataset. Their size is much smaller than that of the average plasmid in complete genomes (∼80 kb), reflecting the split of plasmids across different contigs because of the presence of repeated sequences, *e.g.* IS elements. Hence, we have not attempted to estimate the exact number of plasmids per genome and focus our analysis on the number of genes predicted to be in plasmid contigs. **MGEs (Plasmids + Prophages)**: We found 11,864 gene families specifically related to plasmid elements, 14,188 to prophage elements, and 2,599 shared by both (9% of the MGEs gene families). In complete genomes, prophage and plasmids elements account for half of the pan-genome, of which 1 third were singletons. The large fraction of singletons from MGEs confirms that these elements are extremely diverse and evolved very rapidly, which underlines the difficulty of accurately detecting them and probably leads to their under-estimation in draft genomes. **Loci encoding conjugative or mobilizable elements** were detected with the CONJscan module of MacSyFinder^116^, using protein profiles and definitions following a previous work^54, 117^. 87% of conjugative systems and 75% of putative mobilizable elements were located on contigs predicted as plasmids by Plascope. **Integrons** were identified using IntegronFinder v.1.5 with the –local_max option^57^. 186 integron-integrase (*intI*) were detected with one quarter located at the edges of contigs. We only found one copy per genome. They were often located on very short contigs (20 proteins on average), and five make all the contigs. Most (86%) were located on contigs predicted as plasmid by Plascope, the remaining were on unclassified contigs. Except for the latter, *intI* genes were always located next to ARGs. **IS elements** were identified using ISfinder^56^. Only hits with an e-value lower than 10^-10^, a minimum alignment coverage of 50% and with at least 70% identity were selected, we extracted the IS name of the best hit. Therefore, we identified 47,592 genes encoded for IS elements, among them 43% were located at the edges of contigs (20,329/47,592). They represented 1,006 gene families (∼1% of the pan-genome), of which 41% were singletons. Only 13% were multigenic protein families (*i.e.*, with more than one member in at least one genome). Among them, 9 protein families were found in more than 10 copies in at least one genome, i.e., ISEc1 (10 copies), IS1397 (11), ISSoEn2 (11), IS621 (11), IS2 (15), IS629 (17), IS200C (17) IS1203 (18), and the most extreme case IS1F (107). Very large numbers of ISs, usually a sign of recent proliferation, was restricted to a small number of genomes (Supplementary Dataset1), but this may be an under-estimate caused by the loss of ISs in the assembling process. ISs were often fragmented, characterized by numerous singletons, and six times more frequently present at the edges of contigs than expected by chance. All the results are reported in Supplementary Dataset1.

### Antibiotic resistance genes (ARG)

were detected using 2 curated databases of antibiotic resistance protein: Resfinder v.3.1^118^ and ARG-ANNOT v.3^119^. Therefore, we used BlastP and selected the hits with an e-value lower than 10^-5^, with at least 90% of identity and a minimum alignment coverage of 50%. We found a strong positive correlation between the number of ARGs per genome using each database (pearson’s r=0.97, P<10^-4^). The main difference is the additional detection of three ARGs by ARG-ANNOT, i.e., AmpC2, AmpH, Mfd, which are persistent in Australian dataset and normally do not confer antibiotic resistance in *E. coli*. All the results are reported in Supplementary Dataset1.

### Virulence factors (VF)

were identified using VFDB (downloaded in February 2018, ^120^). The two databases, *i.e.*, VFDB_setA and VFDB_setB were used independently. We used BlastP and selected the hits with an e-value lower than 10^-5^, at least 70% of identity and minimum alignment coverage of 50%. We found 1,332 (vs. 3481) gene families encoding virulence factors with the setA (vs. setB). In spite of these differences, we found qualitatively similar conclusion with the 2 sets because they are very correlated (pearson’s r=0.97, P<10^-4^). All the results are reported in Supplementary Dataset1.

## Acknowledgements

This work was supported by in-house funding from Pasteur Institute and the CNRS (M.T., A.P., JAM.S. and EPC.R.) and was partially supported by grants from the Fondation pour la Recherche Médicale (Equipe FRM 2016, grant number DEQ20161136698, to E.D., and Equipe FRM: EQU201903007835 to EPC.R.) and by an Australian Research Council Linkage Grant (Grant No. LP120100327, D.G., B.V., S.B.). This work used the computational and storage services (TARS cluster) provided by the IT department at Pasteur Institute, Paris.

## Author Contributions

D.G., B.V., S.B., and CL.B. collected, sequenced and assembled the Australian isolates. A.P. annotated the genomes and deposited them into ENA. M.T., E.D., D.G., and EPC.R. designed the research. M.T. managed the project and made most of the computational analyses. JAM.S. performed computational analyses. M.T and EPC.R analyzed the data. M.T. and EPC.R. wrote the manuscript with input from other co-authors.

## Competing Interests Statement

No conflict of interest.

## Data accessibility

Data deposition: Genome assemblies have been deposited into the European Nucleotide Archive (ENA) at EMBL-EBI under accession number PRJEB34791.

This article contains supporting information online at url :

