## Supplementary Information for "Phylogenetic background and habitat drive the genetic diversification of *Escherichia coli*"

### Supplementary Note

Isolates description
Quality control of the genomic sequences
Effect of contig breaks on the estimates of pan-genomes

### Supplementary Tables

Supplementary Table 1: Overall diversity of the three datasets.
Supplementary Table 2: Genetic diversification across phylogroups of Australian dataset.
Supplementary Table 3: The effects of phylogroup and the strains' source on genome size**:** results of the stepwise multiple regression.
Supplementary Table 4: The effects of phylogroup and the strains' source on genome size without MGE**:** results of the stepwise multiple regression.
Supplementary Table 5: Genetic diversification across sources of Australian dataset.
Supplementary Table 6: The effects of phylogroup and the strains' source on MGE content: results of the stepwise multiple regression.

### Supplementary Figures

Supplementary Figure 1: General genomic characteristics of the 1,294 Australian *E. coli* genomes.
Supplementary Figure 2: The large Australian *E. coli* pan-genome.
Supplementary Figure 3: The genus and species phylogenetic trees.
Supplementary Figure 4: Singleton characterization.
Supplementary Figure 5: Comparison of Australian, ECOR and RefSeq datasets.
Supplementary Figure 6: Association between GRR (%, Gene Repertoire Relatedness) and the patristic distance of each pair of genomes.
Supplementary Figure 7: Intra- and Inter-phylogroup genetic diversity.
Supplementary Figure 8: Pan-genomes, Pan-MGE, and rarefied Pan-genomes of each phylogroup and isolation source.
Supplementary Figure 9: Correlation between the different distances and indexes.
Supplementary Figure 10: Gene repertoire relatedness (GRR) within and between phylogroups.
Supplementary Figure 11: Detection of prophages.
Supplementary Figure 12: Detection of plasmid elements.
Supplementary Figure 13: General genomic characteristics of the mobilome.
Supplementary Figure 14: Contribution of MGEs to genome size variation.
Supplementary Figure 15: Distribution of gene families related to MGEs across phylogroups and sources.
Supplementary Figure 16: Network of recent co-occurence of gains (co-gains) of MGE genes within and between phylogroups.

Supplementary Figure 17: Genome size and MGE content according to sources within each phylogroup.

Supplementary Figure 18: Association of integrons and ARGs with human (or domesticated animals).

Supplementary Figure 19: Distribution of VFs and Colicins MGEs across phylogroups and sources.

### Supplementary References

### Supplementary Note

#### Isolates description

The collections of strains from more than 3,300 non-human vertebrate hosts were acquired by sampling a single putative *E. coli* isolate from the feces of each host. They represent diverse species collected from across Australia[1-3]. The mammals sampled were assumed to be healthy hosts. The birds sampled represented a mix of presumably healthy birds collected from the wild, and native birds arriving at veterinarian clinics and wildlife rehabilitation centers. The phylogroup membership of these isolates was determined using the Clermont method[4]. The subset of strains selected for whole genome sequencing (WGS) was chosen to represent the diversity present in each phylogroup.

*E. coli* isolates were recovered from 306 samples of poultry meat purchased from retail outlets in the Australian Capital Territory[5]. Multiple isolates were collected per meat sample, characterized using REP-PCR, and assigned to phylogroups using the Clermont method. One example of each REP-type was selected for WGS.

Two collections of *E. coli* from more than 1,000 humans living in the Canberra region were aquired in 2002 and 2015 by taking a single isolate from unique urine, faecal or blood samples processed by the Canberra Hospital Microbiology Laboratory[6,7]. A subset of strains were selected to represent the diversity present in each phylogroup. Another collection of *E. coli* from 69 humans living in the Canberra region was created by sampling up to 20 *E. coli* isolates from biopsies taken from up to five locations in the lower gut[8]. The isolates were characterised using REP-PCR and assigned to phylogroups using the Clermont method. One example of each REP-type present in a host was selected for WGS.

A collection of *E. coli* isolated from over 800 water samples collected from the Gold Coast region of Queensland, the Sydney region of New South Wales, and the Australian Capital Territory was created by taking up to 20 isolates from each sample[9]. The isolates were characterised using REP-PCR and assigned to phylgroups. For WGS, a subset of isolates was chosen to represent the diversity present in each phylogroup.

### Quality control of the genomic sequences

To assess the completeness of the draft genomes of the Australian and Outgroup datasets, we (i) computed the minimum number of contigs necessary to cover 90% of the genome (L90 value), (ii) the total number of contigs of each genome, and (iii) checked for the presence of known *E. coli* essential genes. ***L90 value***. We computed the L90 value for all draft genomes. A high L90 value means that the genome is scattered into many contigs. Thus, may be caused by abundant repeats, such as insertion sequences (ISs), or poor sequence quality. The distribution of L90 values showed a clear gap between 316 and 385. Therefore, we decided to put the limit at L90=316, and found six genomes with a higher value from this study. ***Number of contigs*****.** We also computed the number of contigs for all draft genomes. The resulting distribution showed a clear gap between 796 and 947 contigs. Therefore, we decided to put the limit at 796 contigs, and found five genomes with a higher value. **E. coli *essential genes*.** We retrieved 296 essential *E. coli* K12-MG1655 genes from the database DEG 10[10] (Supplementary Dataset1). Among them, we excluded 14 genes coding for poison-antidote systems like TA and RMS, or related to phage elements, because they were expected not to be present in all genomes. Homologous genes (>80% identity) were identified in all *Escherichia* genomes (from Australian, RefSeq and Outgroup datasets) using usearch v.11 (option –usearch_global; id=0.8)[11]. The 282 remaining essential genes were all persistent (present in at least 99% of genomes) among the RefSeq dataset. However, some complete genomes were missing up to 6 essential genes. We found that 77% of draft genomes contained all essential genes, 17% were missing one gene, and only four genomes more than 6 essential genes. The latter were removed from further analysis. The results of these 3 independent tests were reported in Supplementary Dataset1. In summary, 10 Australian *E. coli* draft genomes and one outgroup genome failed at least one of these tests and were removed from further analysis, leading to a final dataset of 1,294 Australian *E. coli* genomes and 87 outgroup genomes.

### Effect of contig breaks on the estimates of pan-genomes

The genomes are not completely assembled, which may lead to the observation of spurious gene fragments and gene calling errors. To test the impact of this on the observed genetic diversity, we carried out four tests (Supplementary Fig. 3). First, we compared the average gene size of the three family categories, and found that singletons were almost half the size of persistent (and accessory) genes. The observation that many singletons are small suggests that they are partial genes. Second, to test if singletons were all at contig borders, we computed the fraction of each pan-genome category located at the edges of the contigs. As expected, singletons were largely over-represented (6 times more) at these positions (and those in this location were even smaller). Nevertheless, only 3% of the edges of contigs correspond to singletons, showing that contig breaks only rarely lead to singletons. The low correlation observed between the number of singletons and the number of contigs supports this result (spearman's rho= 0.27, $P<10^{-4}$). Third, we searched for sequence similarities between singletons and the other two categories using Blast+ v.2.6.0[12], and found that 39% of them showed extensive sequence similarity (at least 80% of identity) to larger proteins from the accessory or persistent categories. Finally, we tested if our genomes produced very different results when compared with completely assembled genomes (RefSeq). The comparison of the number of singletons and persistent genes in the rarefied Australian and RefSeq datasets (see Methods), showed that the number of persitent gene families was higher in our dataset. Singletons were 30% more numerous in the Australian genomes, but still represented one third (35%) of the pan-genome of RefSeq genomes. In summary, there is probably an over-prediction of singletons in the Australian dataset (probably one third of the singletons are parts of CDS), but this has a small impact on the definitions of accessory and persistent gene datasets. Together, these two categories make 42,000 different gene families, which remains an impressive number for a single bacterial species. The average number of singletons per genome in each phylogroup of the Australian dataset were not significantly different (Wilcoxon tests, $P>0.05$). Hence, this over-prediction is unlikely to affect most of our results.

### Supplementary Tables

#### Supplementary Table 1: Overall diversity of the three datasets.

| | | Datasets | | |
|---|---|---|---|---|
| | | Australian | RefSeq | ECOR |
| Number of genomes | # | 1294 | 370 | 72 |
| Min-Max Genome Size (GS) | Mb | 4.20-6.02 | 3.98-6.02 | 4.50-5.59 |
| ΔGS | Mb | 1.82 | 2.04 | 1.09 |
| Mean GS | Mb (Std Dev) | 5.02 (0.27) | 5.15 (0.32) | 4.97 (0.26) |
| GC% | % (Std Dev) | 50.64 (0.14) | 50.66 (0.15) | 50.62 (0.14) |
| Gene density % | % (Std Dev) | 87.24 (0.79) | 86.40 (1.36) | 87.55 (0.57) |
| Sequence Type ST | Richness (NA*) | 442 (38) | 135 (16) | 45 (1) |
| | α diversity** | 7.53 | 6.10 | 5.08 |
| H-type | Richness (NA*) | 46 (15) | 37 (10) | 26 |
| | α diversity** | 4.73 | 4.37 | 4.27 |
| O-group | Richness (NA*) | 142 (563) | 96 (24) | 42 (6) |
| | α diversity** | 5.91 | 5.50 | 5.03 |
| O :H serotype | Richness (NA*) | 311 (568) | 175 (32) | 58 (6) |
| | α diversity** | 7.53 | 6.41 | 5.55 |
| Nucleotide diversity Pi | Mean *** (Std Dev) | 0.01078 (0.0078) | 0.01157 (0.0084) | 0.01089 (0.0081) |
| Average-genome | # of proteins | 4683 | 4965 | 4635 |
| Pan-genome | # of protein families**** | 75890 | 33041 | 18584 |
| Persistent-genome (99%) | | 2486 | 2157 | 2494 |
| Core-genome (100%) | | 295 | 667 | 2494 |

* number of untypable genomes (NA)
** Shannon index
*** average of nucleotide diversity ($\pi$) of 112 core gene families in each dataset (ECOR, RefSeq, Australian)
**** from matrix of presence/absence of gene families : gene amplifications were not taken into account

### Supplementary Table 2: Genetic diversification across phylogroups of Australian dataset

| | | Australian dataset | | | | | | | |
|---|---|---|---|---|---|---|---|---|---|
| | | A | B1 (+C) | E | D | F | G | B2 | ALL |
| Genome | # | 312 | 291 (+18) | 61 | 185 | 71 | 33 | 323 | 1294 |
| Min-Max GS | Mb | 4.42-5.81 | 4.20-5.52 | 4.80-5.74 | 4.74-6.02 | 4.77-6.00 | 4.62-5.31 | 4.51-5.51 | 4.20-6.02 |
| Δ GS | Mb | 1.39 | 1.32 | 0.95 | 1.27 | 1.24 | 0.69 | 1.00 | 1.82 |
| Mean GS [1] | Mb (Std Dev) | 4.92 (0.32) | 4.89 (0.19) | 5.22 (0.23) | 5.15 (0.27) | 5.22 (0.26) | 5.07 (0.19) | 5.06 (0.19) | 5.02 (0.27) |
| Proteome [1] | Mean # of genes (Std Dev) | 4611 (347) | 4585 (208) | 4811 (260) | 4770 (317) | 4843 (276) | 4737 (203) | 4730 (218) | 4683 (286) |
| Proteome woMGE [1] | | 4259 (211) | 4237 (122) | 4448 (188) | 4379 (179) | 4450 (166) | 4328 (105) | 4325 (112) | 4309 (174) |
| MGE [1] | | 352 (179) | 349 (134) | 364 (126) | 391 (177) | 393 (142) | 409 (120) | 405 (151) | 375 (158) |
| Sequence Type ST | Richness (NA*) | 92 (13) | 145 (11) | 29 | 57 (7) | 20(2) | 3 | 96 (5) | 442 (38) |
| | α diversity(**) | 4.97 | 6.55 | 3.74 | 5.01 | 3.41 | 0.87 | 4.81 | 7.53 |
| H-type | Richness (NA*) | 31 (8) | 28 | 19 | 23 (1) | 11 | 4 | 16 (6) | 46 (15) |
| | α diversity(**) | 3.98 | 3.94 | 3.53 | 3.61 | 2.58 | 1.16 | 2.93 | 4.73 |
| O-group | Richness (NA*) | 53 (166) | 65 (131) | 23 (26) | 26 (90) | 10 (37) | 6 (20) | 37 (93) | 142 (563) |
| | α diversity(**) | 4.99 | 5.17 | 4.14 | 3.48 | 2.67 | 2.41 | 4.05 | 5.91 |
| O:H serotype | Richness (NA*) | 143 (169) | 178 (131) | 35 (26) | 95 (90) | 34 (37) | 13 (20) | 228 (95) | 311 (568) |
| | α diversity(**) | 5.70 | 6.35 | 4.19 | 4.52 | 3.67 | 2.93 | 5.06 | 7.53 |
| Nucleotide diversity Pi[1] | Mean **** | 0.00425 | 0.00390 | 0.00616 | 0.00749 | 0.00680 | 0.00170 | 0.00451 | 0.01078 |
| | Std Dev | 0.00401 | 0.00449 | 0.00525 | 0.00547 | 0.00629 | 0.00251 | 0.00366 | (0.0078) |
| Average-genome [1] | # of gene families*** | 4510 | 4505 | 4701 | 4669 | 4745 | 4619 | 4642 | 4589 |
| Pan-genome | | 36712 | 32964 | 15097 | 24724 | 14536 | 9435 | 27766 | 75890 |
| α heaps law | | 0.43 | 0.48 | 0.43 | 0.57 | 0.66 | 0.68 | 0.48 | 0.46 |
| Persistent-genome | | 2475 | 2833 | 2714 | 2840 | 2636 | 3437 | 3069 | 2486 |
| Pan/Average | | 8 | 7 | 3 | 5 | 3 | 2 | 6 | 17 |
| Persistent/Average % | | 55 | 63 | 58 | 61 | 56 | 74 | 66 | 54 |
| MGE | | 14011 | 13971 | 4759 | 9552 | 4868 | 2755 | 10161 | 28651 |
| MGE/Pan % | | 38 | 42 | 32 | 39 | 33 | 29 | 37 | 38 |
| Phage-related [1] | Mean # of genes (Std Dev) | 199 (94) | 223 (91) | 222 (69) | 247 (110) | 228 (89) | 231 (70) | 250 (107) | 228 (99) |
| Plasmid-related [1] | | 166 (135) | 133 (97) | 143 (92) | 164 (133) | 194 (103) | 191 (101) | 154 (94) | 157 (113) |
| IS-related [1] | | 46 (21) | 29 (15) | 61 (39) | 36 (22) | 41 (20) | 44 (18) | 34 (14) | 37 (19) |
| VFA [2] | | 95 (15) | 102 (11) | 115 (8) | 116 (10) | 113 (9) | 113 (10) | 124 (11) | 109 (16) |
| VFB [2] | | 199 (29) | 225 (21) | 245 (15) | 242 (17) | 237 (16) | 219 (12) | 242 (25) | 227 (29) |
| ARG Resfinder [2] | | 2.3 (2.2) | 1.9 (1.8) | 2.1 (1.3) | 1.8 (1.9) | 4.8 (3.8) | 3.1 (2.6) | 2.2 (2.0) | 2.2 (1.2) |
| ARG Arganot [2] | | 5.4 (2.5) | 5.0 (2.0) | 5.3 (1.5) | 4.5 (2.2) | 7.9 (3.9) | 6.2 (2.8) | 4.2 (2.1) | 5.0 (2.5) |
| *int1+* [3] | % of genomes | 16% | 10% | 8% | 6% | 39% | 30% | 16% | 14% |
| CONJ+[2] | | 39% | 54% | 72% | 53% | 45% | 58% | 66% | 54% |
| Rarefied-PanGenome [1,4] N=50 | Mean # of gene families (Std Dev) | 15935 (619) | 14446 (525) | 13890 (322) | 14861 (486) | 12784 (373) | NA [4] | 13246 (551) | |
| Rarefied-Persistent [1,4] N=50 | | 2367 (80) | 2699 (132) | 2831 (41) | 2915 (77) | 2838 (41) | NA [4] | 2954 (77) | |
| Rarefied-MGE [1,4] N=50 | | 6477 (335) | 6369 (314) | 4338 (140) | 5921 (248) | 4213 (161) | NA [4] | 5046 (306) | |

* number of untypable genomes (NA)

** Shannon index

*** matrix of presence/absence of gene families : gene amplifications were not taken into account

**** mean of 112 core genes (the same as those of the 3 datasets : ECOR, RefSeq and Australian)

(1) standard ANOM test

(2) non-parametric ANOM test

(3) ANOM for proportions

(4) rarefied datasets were computed from 1000 combinations of 50 distinct genomes, while G group contains only 33 genomes.

**Supplementary Table 3: The effects of phylogroup and the strains' source on genome size (Mb): results of the stepwise multiple regression.** Stepwise regression is an approach to selecting a subset of parameters (among the strains' source and phylogroup) for a regression model. In forward selection, terms are entered into the model and most significant terms are added until all of the terms are significant. We used the minimum Bayesian Information Criterion to choose the best model. The Stepwise regression report (1) shows the statistics of the best model. As each step is taken, the Step History report (2) records the effect of adding a term to the model, and shows the order in which the terms entered the model and the statistics for each model. The Current Estimates report (3) indicates whether a term is currently in the best model and shows the statistics of each term for this model.

**1- STEPWISE REGRESSION REPORT**

| Y | SSE (1) | DFE (2) | RMSE (3) | RSquare (4) | Rsquare Adj (5) | Cp (6) | p (7) | AICc (8) | BIC (9) |
|---|---|---|---|---|---|---|---|---|---|
| **Genome_size** | **5.9028E+13** | **1269** | **215674.8678** | **0.360** | **0.357** | **20.7448** | **6** | **34944.26** | **34980.23** |

(1) SSE : Sum of squared errors for the current model

(2) DFE : Error degrees of freedom for the current model

(3) RMSE : Root mean square error (residual) for the current model

(4) RSquare : Proportion of the variation in the response that can be attributed to terms in the model rather than to random error.

(5) RSquare Adj: Adjusts R2 to make it more comparable over models with different numbers of parameters by using the degrees of freedom in its computation.
The adjusted R2 is useful in stepwise procedure because you are looking at many different models and want to adjust for the number of terms in the model.

(6) Cp : Mallow's Cp criterion for selecting a model.

(7) p : Number of parameters in the model, including the intercept.

(8) AICc : Corrected Akaike's Information Criterion

(9) BIC : Bayesian Information Criterion

**2-STEP HISTORY REPORT**

| Y | Step | | Parameter (12) | ACTION (10) | "Sig Prob" (11) | RSquare (4) | Cp (6) | p (7) | AICc (8) | BIC (9) |
|---|---|---|---|---|---|---|---|---|---|---|
| **Genome size (Mb)** | **1** | Source | **Source{Water&BF&HI&MF&HF-HE&PM}** | **Entered** | **1.04698E-54** | **0.174** | **386.276** | **2** | **35261.82** | **35277.25** |
| **Genome size (Mb)** | **2** | Phylogroup | **Phylogroup{B1&A-B2&G&D&F&E}** | **Entered** | **3.09628E-50** | **0.306** | **122.628** | **3** | **35041.09** | **35061.66** |
| **Genome size (Mb)** | **3** | Phylogroup | **Phylogroup{B2&G-D&F&E}** | **Entered** | **5.57312E-12** | **0.332** | **73.587** | **4** | **34995.46** | **35021.17** |
| **Genome size (Mb)** | **4** | Source | **Source{Water&BF-HI&MF&HF}** | **Entered** | **4.17722E-08** | **0.347** | **44.211** | **5** | **34967.29** | **34998.12** |
| **Genome size (Mb)** | **5** | Source | **Source{HE-PM}** | **Entered** | **5.98408E-07** | **0.360** | **20.745** | **6** | **34944.26** | **34980.23** |
| Genome size (Mb) | 6 | Source | Source{HI-MF&HF} | Entered | 0.021480237 | 0.363 | 17.401 | 7 | 34940.97 | 34982.06 |
| Genome size (Mb) | 7 | Phylogroup | Phylogroup{B2-G} | Entered | 0.062954831 | 0.364 | 15.915 | 8 | 34939.52 | 34985.73 |
| Genome size (Mb) | 8 | Source | Source{MF-HF} | Entered | 0.050472158 | 0.366 | 14.067 | 9 | 34937.69 | 34989.03 |
| Genome size (Mb) | 9 | Phylogroup | Phylogroup{B1-A} | Entered | 0.04915779 | 0.368 | 12.183 | 10 | 34935.83 | 34992.28 |
| Genome size (Mb) | 10 | Source | Source{Water-BF} | Entered | 0.080586062 | 0.370 | 11.125 | 11 | 34934.78 | 34996.35 |
| Genome size (Mb) | 11 | Phylogroup | Phylogroup{D-F&E} | Entered | 0.165659919 | 0.371 | 11.202 | 12 | 34934.88 | 35001.56 |
| Genome size (Mb) | 12 | Phylogroup | Phylogroup{F-E} | Entered | 0.653360514 | 0.371 | 13.000 | 13 | 34936.73 | 35008.50 |
| **Genome size (Mb)** | **13** | | **Best model** | **Specific** | | **0.360** | **20.745** | **6** | **34944.26** | **34980.23** |

(10) ACTION : Entered = Indicates whether a term is currently in the model.

(11) "Sig Prob" : The significance level associated with the Wald/Score ChiSq test statistic based on nDF degrees of freedom. The "Sig Prob" is used to determine the next term to be included in the model.

(12) Parameter : Water=freshwater; BF=Bird Faecal; HI=Human Intestinal, MF = Mammal Faecal; HF = Human Faecal ; HE = Human Extra-intestinal; PM = Poultry Meat

**3-CURRENT ESTIMATES REPORT**

| Y | ACTION (10) | | Parameter (12) | Estimate (13) | nDF (14) | SS (15) | "F Ratio" (16) | "Prob>F" (17) | RSquare (4) | %Explained_ Variance |
|---|---|---|---|---|---|---|---|---|---|---|
| Genome_size | Entered | | Intercept | 5047531.05 | 1 | 0 | 0 | 1 | | |
| **Genome_size** | **Entered** | **Source** | **Source{Water&BF&HI&MF&HF-HE&PM}** | **-100800.31** | **3** | **1.7235E+13** | **123.508** | **3.4692E-70** | **0.174** | **48.2** |
| **Genome_size** | **Entered** | **Source** | **Source{Water&BF-HI&MF&HF}** | **-39909.963** | **1** | **1.3263E+12** | **28.513** | **1.1017E-07** | **0.016** | **4.3** |
| Genome_size | | Source | Source{Water-BF} | 0 | 1 | 1.2092E+11 | 2.603 | 0.10691475 | | |
| Genome_size | | Source | Source{HI-MF&HF} | 0 | 1 | 2.4573E+11 | 5.301 | 0.02148024 | | |
| Genome_size | | Source | Source{MF-HF} | 0 | 1 | 1.5761E+11 | 3.395 | 0.06563984 | | |
| **Genome_size** | **Entered** | **Source** | **Source{HE-PM}** | **-62298.379** | **1** | **1.171E+12** | **25.174** | **5.9841E-07** | **0.013** | **3.5** |
| **Genome_size** | **Entered** | **Phylogroup** | **Phylogroup{B1&A-B2&G&D&F&E}** | **-97045.169** | **2** | **1.3348E+13** | **143.482** | **6.6454E-57** | **0.132** | **36.8** |
| Genome_size | | Phylogroup | Phylogroup{B1-A} | 0 | 1 | 1.0314E+11 | 2.219 | 0.13653071 | | |
| **Genome_size** | **Entered** | **Phylogroup** | **Phylogroup{B2&G-D&F&E}** | **-54784.388** | **1** | **1.8827E+12** | **40.475** | **2.7748E-10** | **0.025** | **7.1** |
| Genome_size | | Phylogroup | Phylogroup{B2-G} | 0 | 1 | 1.3688E+11 | 2.947 | 0.08627447 | | |
| Genome_size | | Phylogroup | Phylogroup{D-F&E} | 0 | 1 | 1.523E+11 | 3.280 | 0.07035921 | | |
| Genome_size | | Phylogroup | Phylogroup{F-E} | 0 | 1 | 1.7961E+10 | 0.386 | 0.53455457 | | |

(13) Estimate :The current parameter estimate, which is zero if the effect is not currently in the model

(14) nDF : The number of degrees of freedom for a term. A term has more than one degree of freedom if its entry into a model also forces other terms into the model.

(15) SS : The reduction in the error (residual) sum of squares (SS) if the term is entered into the model or the increase in the error SS if the term is removed from the model.

(16) F ratio: The traditional test statistic to test that the term effect is zero. It is the square of a t-ratio.

(17) Prob>F : The significance level associated with the F statistic.

**Supplementary Table 4: The effects of phylogroup and the strains' source on genome size without MGE (number of genes): results of the stepwise multiple regression.** Same approach described in Supplementary Table 3.

**1- STEPWISE REGRESSION REPORT**

| Y | SSE (1) | DFE (2) | RMSE (3) | RSquare (4) | Rsquare Adj (5) | Cp (6) | p (7) | AICc (8) | BIC (9) |
|---|---|---|---|---|---|---|---|---|---|
| **GenomeSize-woMGE** | **27365270.2** | **1267** | **146.9642** | **0.293** | **0.289** | **7.6848** | **8** | **16353.39** | **16399.61** |

(1) SSE : Sum of squared errors for the current model

(2) DFE : Error degrees of freedom for the current model

(3) RMSE : Root mean square error (residual) for the current model

(4) RSquare : Proportion of the variation in the response that can be attributed to terms in the model rather than to random error.

(5) RSquare Adj: Adjusts R2 to make it more comparable over models with different numbers of parameters by using the degrees of freedom in its computation.
The adjusted R2 is useful in stepwise procedure because you are looking at many different models and want to adjust for the number of terms in the model.

(6) Cp : Mallow's Cp criterion for selecting a model.

(7) p : Number of parameters in the model, including the intercept.

(8) AICc : Corrected Akaike's Information Criterion

(9) BIC : Bayesian Information Criterion

**2-STEP HISTORY REPORT**

| Y | Step | | Parameter (12) | ACTION (10) | "Sig Prob" (11) | RSquare (4) | Cp (6) | p (7) | AICc (8) | BIC (9) |
|---|---|---|---|---|---|---|---|---|---|---|
| **Genome_Size-woMGE** | **1** | Source | **Source{Water&BF&HI&MF&HF-HE&PM}** | **Entered** | **0** | **0.1239** | **298.269** | **2** | **16614.38** | **16629.82** |
| **Genome_Size-woMGE** | **2** | Phylogroup | **Phylogroup{B1&A-B2&G&D&E&F}** | **Entered** | **0** | **0.2232** | **122.315** | **3** | **16462.94** | **16483.51** |
| **Genome_Size-woMGE** | **3** | Phylogroup | **Phylogroup{B2&G&D-E&F}** | **Entered** | **0** | **0.2449** | **85.391** | **4** | **16428.77** | **16454.48** |
| **Genome_Size-woMGE** | **4** | Source | **Source{Water&BF-HI&MF&HF}** | **Entered** | **0** | **0.264** | **53.300** | **5** | **16398.24** | **16429.08** |
| **Genome_Size-woMGE** | **5** | Phylogroup | **Phylogroup{B2&G-D}** | **Entered** | **0** | **0.2795** | **27.573** | **6** | **16373.16** | **16409.13** |
| **Genome_Size-woMGE** | **6** | Source | **Source{HE-PM}** | **Entered** | **0.0001** | **0.2883** | **13.742** | **7** | **16359.45** | **16400.54** |
| **Genome_Size-woMGE** | **7** | Source | **Source{HI-MF&HF}** | **Entered** | **0.0046** | **0.2928** | **7.685** | **8** | **16353.39** | **16399.61** |
| Genome_Size-woMGE | 8 | Phylogroup | Phylogroup{B2-G} | Entered | 0.1664 | 0.2939 | 7.770 | 9 | 16353.49 | 16404.83 |
| Genome_Size-woMGE | 9 | Source | Source{MF-HF} | Entered | 0.1655 | 0.2949 | 7.848 | 10 | 16353.59 | 16410.04 |
| Genome_Size-woMGE | 10 | Phylogroup | Phylogroup{B1-A} | Entered | 0.4617 | 0.2952 | 9.306 | 11 | 16355.08 | 16416.64 |
| Genome_Size-woMGE | 11 | Phylogroup | Phylogroup{E-F} | Entered | 0.5938 | 0.2954 | 11.022 | 12 | 16356.84 | 16423.51 |
| Genome_Size-woMGE | 12 | Source | Source{Water-BF} | Entered | 0.8822 | 0.2954 | 13.000 | 13 | 16358.86 | 16430.64 |
| **Genome_Size-woMGE** | **13** | | **Best model** | **Specific** | | **0.2928** | **7.68483187** | **8** | **16353.3924** | **16399.6064** |

(10) ACTION : Entered = Indicates whether a term is currently in the model.

(11) "Sig Prob" : The significance level associated with the Wald/Score ChiSq test statistic based on nDF degrees of freedom. The "Sig Prob" is used to determine the next term to be included in the model.

(12) Parameter : Water=freshwater; BF=Bird Faecal; HI=Human Intestinal, MF = Mammal Faecal; HF = Human Faecal ; HE = Human Extra-intestinal; PM = Poultry Meat

**3-CURRENT ESTIMATES REPORT**

| Y | ACTION (10) | | Parameter (12) | Estimate (13) | nDF (14) | SS (15) | "F Ratio" (16) | "Prob>F" (17) | RSquare (4) | %Explained_ Variance |
|---|---|---|---|---|---|---|---|---|---|---|
| Genome_Size-woMGE | | | Intercept | 4340.42809 | 1 | 0 | 0 | 1 | | |
| **Genome_Size-woMGE** | **YES** | **Phylogroup** | **Phylogroup{B1&A-B2&G&D&E&F}** | **-70.160057** | **3** | **4945723.63** | **76.328** | **2.19E-45** | **0.099** | **33.9** |
| Genome_Size-woMGE | | Phylogroup | Phylogroup{B1-A} | 0 | 1 | 6030.94294 | 0.279 | 0.59740327 | | |
| **Genome_Size-woMGE** | **YES** | **Phylogroup** | **Phylogroup{B2&G&D-E&F}** | **-42.180222** | **2** | **1294483.42** | **29.967** | **1.92E-13** | **0.022** | **7.4** |
| **Genome_Size-woMGE** | **YES** | **Phylogroup** | **Phylogroup{B2&G-D}** | **-30.200864** | **1** | **420168.543** | **19.454** | **1.1181E-05** | **0.016** | **5.3** |
| Genome_Size-woMGE | | Phylogroup | Phylogroup{B2-G} | 0 | 1 | 41374.5213 | 1.917 | 0.16643077 | | |
| Genome_Size-woMGE | | Phylogroup | Phylogroup{E-F} | 0 | 1 | 7715.72783 | 0.357 | 0.55025406 | | |
| **Genome_Size-woMGE** | **YES** | **Source** | **Source{Water&BF&HI&MF&HF-HE&PM}** | **-54.109454** | **4** | **5455697.05** | **63.149** | **1.02E-48** | **0.124** | **42.3** |
| **Genome_Size-woMGE** | **YES** | **Source** | **Source{Water&BF-HI&MF&HF}** | **-28.053928** | **2** | **911401.279** | **21.099** | **9.69E-10** | **0.019** | **6.5** |
| Genome_Size-woMGE | | Source | Source{Water-BF} | 0 | 1 | 390.086485 | 0.018 | 0.89315625 | | |
| **Genome_Size-woMGE** | **YES** | **Source** | **Source{HI-MF&HF}** | **-21.20986** | **1** | **174066.07** | **8.059** | **0.00459964** | **0.005** | **1.5** |
| Genome_Size-woMGE | | Source | Source{MF-HF} | 0 | 1 | 36616.8256 | 1.696 | 0.19301278 | | |
| **Genome_Size-woMGE** | **YES** | **Source** | **Source{HE-PM}** | **-33.951573** | **1** | **347262.466** | **16.078** | **6.4332E-05** | **0.009** | **3.0** |

(13) Estimate :The current parameter estimate, which is zero if the effect is not currently in the model

(14) nDF : The number of degrees of freedom for a term. A term has more than one degree of freedom if its entry into a model also forces other terms into the model.

(15) SS : The reduction in the error (residual) sum of squares (SS) if the term is entered into the model or the increase in the error SS if the term is removed from the model.

(16) F ratio: The traditional test statistic to test that the term effect is zero. It is the square of a t-ratio.

(17) Prob>F : The significance level associated with the F statistic.

### Supplementary Table 5: Genetic diversification across sources of Australian dataset.

| | | Australian dataset | | | | | | | |
|---|---|---|---|---|---|---|---|---|---|
| | | BF | PM | HE | HI | HF | MF | Water | ALL |
| Genome | # | 195 | 283 | 112 | 172 | 93 | 135 | 285 | 1294 |
| Min-Max GS | Mb | 4.45-6.02 | 4.6-6.00 | 4.62-5.46 | 4.63-5.52 | 4.54-5.55 | 4.42-5.74 | 4.20-5.64 | 4.20-6.02 |
| Δ GS | Mb | 1.57 | 1.44 | 0.84 | 0.89 | 1.01 | 1.32 | 1.44 | 1.82 |
| Mean GS [(1)] | Mb (Std Dev) | 4.93 (0.24) | 5.22 (0.24) | 5.10 (0.18) | 4.99 (0.19) | 5.02 (0.22) | 5.01 (0.22) | 4.87 (0.24) | 5.02 (0.27) |
| Proteome [(1)] | Mean # of genes (Std Dev) | 4578 (248) | 4895 (311) | 4794 (199) | 4643 (194) | 4708 (247) | 4675 (249) | 4525 (240) | 4683 (286) |
| Proteome woMGE [(1)] | | 4243 (154) | 4419 (214) | 4353 (108) | 4291 (124) | 4321 (131) | 4321 (158) | 4229 (144) | 4309 (174) |
| MGE[(1)] | | 334 (144) | 475 (151) | 441 (135) | 352 (123) | 387 (148) | 355 (158) | 296 (144) | 375 (158) |
| Sequence Type ST | Richness (NA*) | 126 (5) | 65 (11) | 39 (4) | 67 (1) | 41 (1) | 91 (5) | 172 (10) | 442 (38) |
| | α diversity(**) | 6.61 | 5.29 | 3.66 | 5.45 | 4.40 | 6.29 | 6.83 | 7.53 |
| H-type | Richness (NA*) | 37 (1) | 33 | 17 | 30 (5) | 21 (3) | 31 (2) | 39 (1) | 46 (15) |
| | α diversity(**) | 4.87 | 4.32 | 2.93 | 4.16 | 3.22 | 4.59 | 4.74 | 4.73 |
| O-group | Richness (NA*) | 49 (101) | 47 (163) | 23 (24) | 31 (73) | 25 (46) | 45 (45) | 75 (105) | 142 (563) |
| | α diversity(**) | 5.20 | 5.00 | 3.43 | 4.65 | 4.36 | 4.88 | 5.20 | 5.91 |
| O:H serotype | Richness (NA*) | 78 (101) | 66 (163) | 30 (25) | 46 (77) | 30 (46) | 65 (46) | 108 (106) | 311 (568) |
| | α diversity(**) | 6.16 | 5.61 | 4.12 | 5.27 | 4.59 | 5.85 | 6.22 | 7.53 |
| Average-genome [(1)] | # of gene families*** | 4504 | 4764 | 4685 | 4562 | 4602 | 4591 | 4454 | 4589 |
| Pan-genome | | 30150 | 28576 | 19253 | 22284 | 20885 | 28391 | 34687 | 75890 |
| α heaps law | | 0.44 | 0.54 | 0.59 | 0.64 | 0.57 | 0.42 | 0.49 | 0.46 |
| Persistent-genome | | 2379 | 2474 | 2801 | 2572 | 2020 [(4)] | 2599 | 2515 | 2486 |
| Pan/Average | | 7 | 6 | 4 | 5 | 5 | 6 | 8 | 17 |
| Persistent/Average % | | 53 | 52 | 60 | 56 | 44 | 57 | 56 | 54 |
| MGE | | 11049 | 10382 | 6317 | 7668 | 6956 | 9802 | 12568 | 28651 |
| MGE/Pan % | | 37 | 36 | 33 | 34 | 33 | 35 | 36 | 38 |
| Phage-related [(1)] | Mean # of genes (Std Dev) | 213 (90) | 254 (95) | 275 (102) | 206 (79) | 220 (102) | 235 (111) | 208 (99) | 228 (99) |
| Plasmid-related [(1)] | | 141 (102 | 245 (120) | 168 (75) | 145 (96) | 164 (103) | 124 (98) | 97 (90) | 157 (113) |
| IS-related [(1)] | | 29 (14) | 54 (19) | 41 (11) | 33 (17) | 43 (20) | 31 (19) | 26 (13) | 37 (19) |
| VFA [(2)] | | 105 (15) | 108 (14) | 118 (17) | 116 (17) | 113 (20) | 112 (16) | 103 (14) | 109 (16) |
| VFB [(2)] | | 219 (29) | 230 (26) | 244 (33) | 233 (30) | 227 (36) | 229 (26) | 218 (25) | 227 (29) |
| ARG Resfinder [(2)] | | 2.1 (2.1) | 2.9 (2.0) | 3.7 (3.2) | 2.7 (2.8) | 2.6 (2.5) | 1.2 (0.8) | 1.3 (1.2) | 2.2 (1.2) |
| ARG Argannot [(2)] | | 5.1 (2.7) | 5.8 (2.3) | 6.2 (3.3) | 5.3 (3.1) | 5.3 (2.7) | 3.8 (1.1) | 4.1 (1.4) | 5.0 (2.5) |
| *int1+* [(3)] | % of genomes | 7% | 24% | 40% | 20% | 20% | <1% | 2% | 14% |
| CONJ+[(2)] | | 48% | 66% | 64% | 57% | 55% | 52% | 41% | 54% |
| Rarefied-PanGenome [(1,5)] N=50 | Mean # of gene families (Std Dev) | 16141 (551) | 15355 (502) | 13837 (520) | 14423 (483) | 16099 (425) | 17800 (557) | 15497 (580) | |
| Rarefied-Persistent [(1,5)] N=50 | | 2486 (71) | 2488 (86) | 2601 (61) | 2628 (64) | 2390 (60) | 2486 (79) | 2506 (91) | |

* number of untypable genomes (NA)
** Shannon index
*** matrix of presence/absence of gene families : gene amplifications were not taken into account
**** mean of 112 core genes (the same as those of the 3 datasets : ECOR, RefSeq and Australian)
(1) standard ANOM test
(2) non-parametric ANOM test
(3) ANOM for proportions
(4) # of genomes <100; hence persistent genome = core genome in this case.
(5) rarefied datasets were computed from 1000 combinations of 50 distinct genomes.

### Supplementary Table 6: The effects of phylogroup and the strains' source on MGE content (number of genes): results of the stepwise multiple regression.

Same approach described in Supplementary Table 3.

#### 1- STEPWISE REGRESSION REPORT

| Y | SSE (1) | DFE (2) | RMSE (3) | RSquare (4) | Rsquare Adj (5) | Cp (6) | p (7) | AICc (8) | BIC (9) |
|---|---|---|---|---|---|---|---|---|---|
| **MGE content** | **25850780.4** | **1269** | **142.7270** | **0.190** | **0.187** | **10.577** | **6** | **16276.75** | **16312.71** |

(1) SSE : Sum of squared errors for the current model

(2) DFE : Error degrees of freedom for the current model

(3) RMSE : Root mean square error (residual) for the current model

(4) RSquare : Proportion of the variation in the response that can be attributed to terms in the model rather than to random error.

(5) RSquare Adj: Adjusts R2 to make it more comparable over models with different numbers of parameters by using the degrees of freedom in its computation.

The adjusted R2 is useful in stepwise procedure because you are looking at many different models and want to adjust for the number of terms in the model.

(6) Cp : Mallow's Cp criterion for selecting a model.

(7) p : Number of parameters in the model, including the intercept.

(8) AICc : Corrected Akaike's Information Criterion

(9) BIC : Bayesian Information Criterion

#### 2-STEP HISTORY REPORT

| Y | Step | | Parameter (12) | ACTION (10) | "Sig Prob" (11) | RSquare (4) | Cp (6) | p (7) | AICc (8) | BIC (9) |
|---|---|---|---|---|---|---|---|---|---|---|
| **MGE content** | **1** | Source | **Source{Water&BF&HI&MF&HF-HE&PM}** | **Entered** | **0** | **0.147** | **69.924** | **2** | **16334.38** | **16349.81** |
| **MGE content** | **2** | Phylogroup | **Phylogroup{B1&A&E-D&F&B2&G}** | **Entered** | **0** | **0.166** | **41.642** | **3** | **16307.27** | **16327.84** |
| **MGE content** | **3** | Source | **Source{Water-BF&HI&MF&HF}** | **Entered** | **0** | **0.179** | **23.314** | **4** | **16289.35** | **16315.06** |
| **MGE content** | **4** | Source | **Source{HE-PM}** | **Entered** | **0.003** | **0.185** | **16.396** | **5** | **16282.53** | **16313.37** |
| **MGE content** | **5** | Source | **Source{BF&HI&MF-HF}** | **Entered** | **0.0053** | **0.190** | **10.577** | **6** | **16276.75** | **16312.71** |
| MGE content | 6 | Phylogroup | Phylogroup{B1-A} | Entered | 0.027 | 0.195 | 7.337 | 8 | 16273.53 | 16319.75 |
| MGE content | 7 | Phylogroup | Phylogroup{B2-G} | Entered | 0.2778 | 0.196 | 8.775 | 10 | 16275.02 | 16331.47 |
| MGE content | 8 | Phylogroup | Phylogroup{D-F} | Entered | 0.2716 | 0.197 | 9.567 | 11 | 16275.84 | 16337.40 |
| MGE content | 9 | Source | Source{BF-HI&MF} | Entered | 0.6634 | 0.197 | 11.377 | 12 | 16277.69 | 16344.36 |
| MGE content | 10 | Source | Source{HI-MF} | Entered | 0.5392 | 0.197 | 13.000 | 13 | 16279.35 | 16351.13 |
| **MGE content** | **11** | | **Best model** | **Specific** | | **0.190** | **10.577** | **6** | **16276.75** | **16312.71** |

(10) ACTION : Entered = Indicates whether a term is currently in the model.

(11) "Sig Prob" : The significance level associated with the Wald/Score ChiSq test statistic based on nDF degrees of freedom. The "Sig Prob" is used to determine the next term to be included in the model.

(12) Parameter : Water=freshwater; BF=Bird Faecal; HI=Human Intestinal, MF = Mammal Faecal; HF = Human Faecal ; HE = Human Extra-intestinal; PM = Poultry Meat

#### 3-CURRENT ESTIMATES REPORT

| Y | ACTION (10) | | Parameter (12) | Estimate (13) | nDF (14) | SS (15) | "F Ratio" (16) | "Prob>F" (17) | RSquare (4) | %Explained_ Variance |
|---|---|---|---|---|---|---|---|---|---|---|
| MGE content | | | Intercept | 395.206327 | 1 | 0 | 0 | 1 | | |
| **MGE content** | **Entered** | Phylogroup | **Phylogroup{B1&A&E-D&F&B2&G}** | **-21.028331** | **1** | **524366.694** | **25.741** | **4.49E-07** | **0.019** | **10.2** |
| MGE content | | Phylogroup | Phylogroup{B1&A-E} | 0 | 1 | 8643.56979 | 0.424 | 0.51500869 | | |
| MGE content | | Phylogroup | Phylogroup{B1-A} | 0 | 2 | 146965.821 | 3.622 | 0.02700251 | | |
| MGE content | | Phylogroup | Phylogroup{D&F-B2&G} | 0 | 1 | 10004.7586 | 0.491 | 0.48364091 | | |
| MGE content | | Phylogroup | Phylogroup{D-F} | 0 | 2 | 32196.297 | 0.79 | 0.45407397 | | |
| MGE content | | Phylogroup | Phylogroup{B2-G} | 0 | 2 | 51206.4187 | 1.257 | 0.28475944 | | |
| **MGE content** | **Entered** | Source | **Source{Water&BF&HI&MF&HF-HE&PM}** | **-59.923462** | **4** | **5353107.24** | **65.695** | **1.51E-50** | **0.147** | **77.4** |
| **MGE content** | **Entered** | Source | **Source{Water-BF&HI&MF&HF}** | **-30.671433** | **2** | **552870.98** | **13.57** | **1.47E-06** | **0.013** | **6.8** |
| **MGE content** | **Entered** | Source | **Source{BF&HI&MF-HF}** | **-22.50959** | **1** | **158694.835** | **7.79** | **0.00533165** | **0.005** | **2.6** |
| MGE content | | Source | Source{BF-HI&MF} | 0 | 1 | 11332.9911 | 0.556 | 0.45595975 | | |
| MGE content | | Source | Source{HI-MF} | 0 | 2 | 17104.9559 | 0.419 | 0.65749817 | | |
| **MGE content** | **Entered** | Source | **Source{HE-PM}** | **-24.267943** | **1** | **183640.521** | **9.015** | **0.0027304** | **0.006** | **3.0** |

(13) Estimate :The current parameter estimate, which is zero if the effect is not currently in the model

(14) nDF : The number of degrees of freedom for a term. A term has more than one degree of freedom if its entry into a model also forces other terms into the model.

(15) SS : The reduction in the error (residual) sum of squares (SS) if the term is entered into the model or the increase in the error SS if the term is removed from the model.

(16) F ratio: The traditional test statistic to test that the term effect is zero. It is the square of a t-ratio.

(17) Prob>F : The significance level associated with the F statistic.

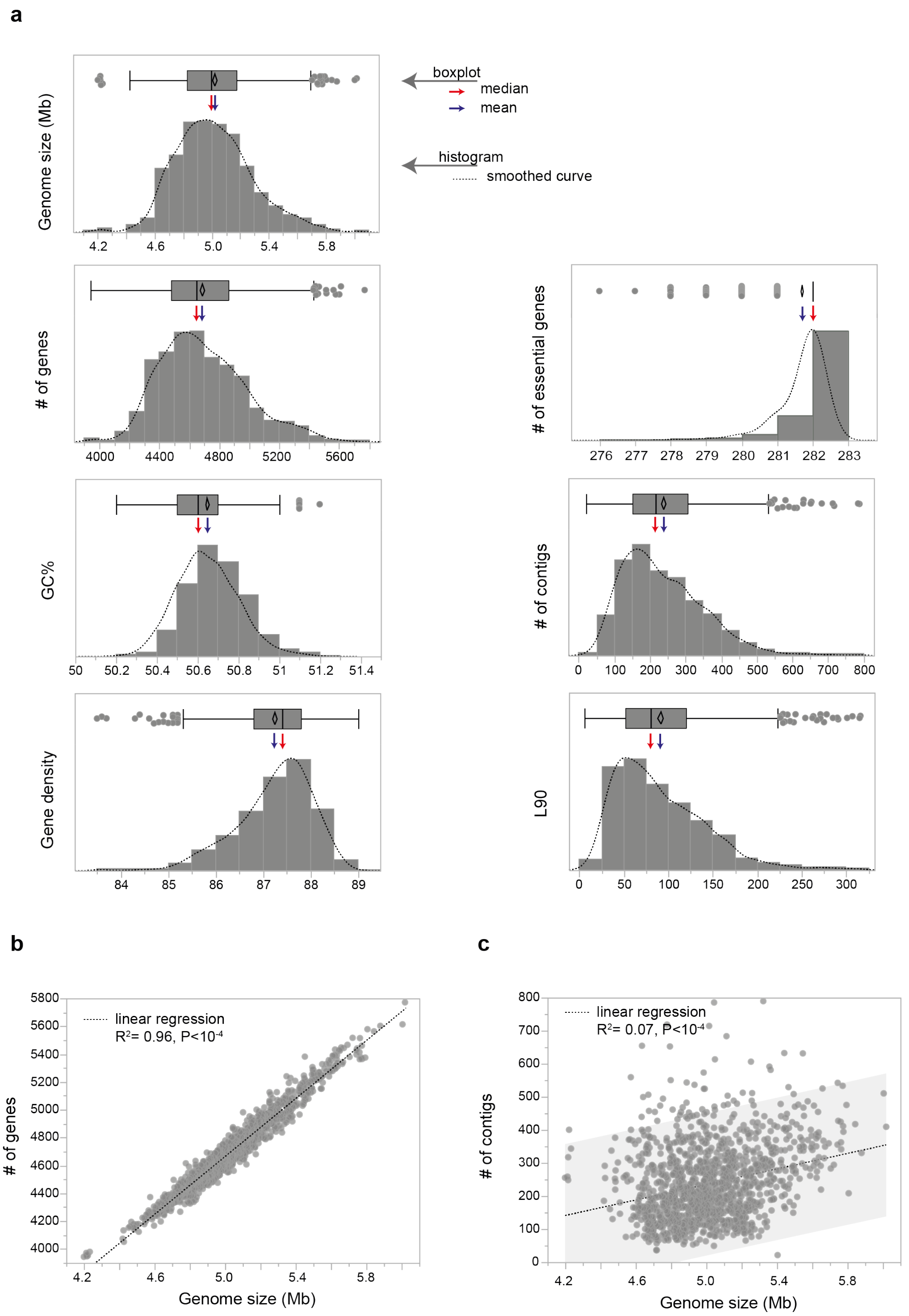


**Supplementary Figure 1**: **General genomic characteristics of the 1,294 Australian *E. coli* genomes. (a)** Histogram and boxplot of genomic features, *i.e.*, the genome size (Mb), the number (#) of genes encoding proteins, the GC content (GC%), the gene density, the number of essential genes, the number of contigs and the L90 (Methods). For each case, the dash line corresponds to the smoothed curve, the red arrow to the median and the blue arrow to the average of each distribution. **(b)** Strong positive correlation between the genome size and the number of genes (spearman's rho= 0.98, P<$10^{-4}$). **(c)** Weak positive correlation between the genome size and the number of contigs (spearman's rho= 0.23, P<$10^{-4}$). The genomes with the greatest number of contigs were not necessarily the largest. Linear regression (dash line) and statistics were reported.

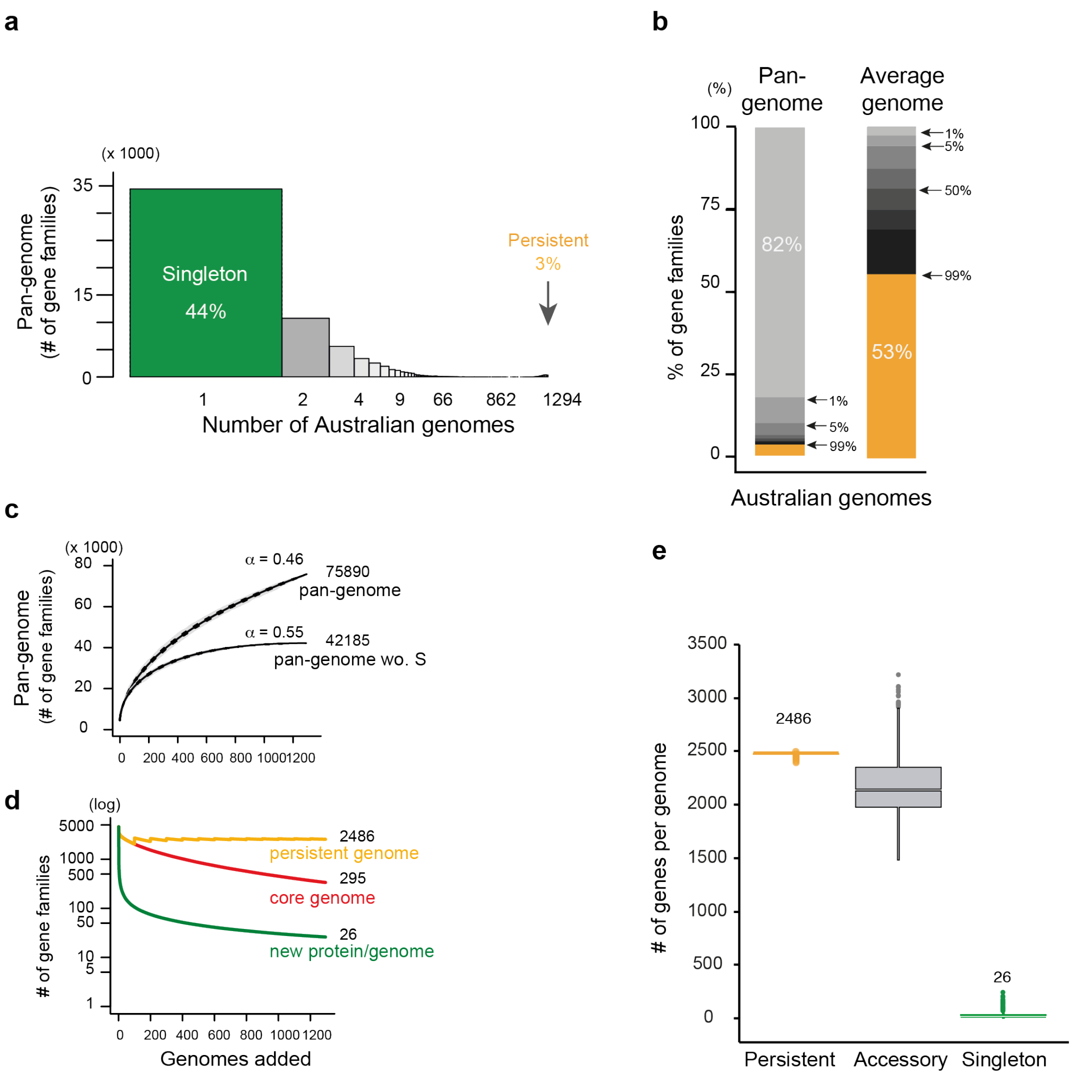


**Supplementary Figure 2: The large Australian *E. coli* pan-genome.** **(a)** Number of gene families according to their occurrence in genomes. Singletons (in green), *i.e.*, genes present in a single genome, represent 44% of the pan-genome. Persistent gene families (in gold), *i.e.*, present in at least 99% of genomes, represent only 3% of the pan-genome. **(b)** Fraction of gene families (%) according to their frequency among the pan-genome and the average genome. Frequencies were represented by a color code ranging from light grey (present in less than 1% of genomes) to black (up to 99%), persistent genes (>99%) were represented in gold. 82% of the gene families are rare, *i.e.*, present in less than 1% of genomes including the 33,705 singletons. Persistent gene families represent 53% of the average genome, while singletons less than 1%. **(c)** Rarefaction curve of the full pan-genome and of the pan-genome after removing the 33,705 singletons (wo. S). In each case, we used 1,000 permutations (genomes orderings) and then averaged the results. The *alpha* (inferred using the heaps' law model) is lower than 1 in both, indicating that the pan-genome is open in both. **(d)** Rarefaction curve of the persistent genome (in gold) and of the core genome (in red), *i.e.*, the cumulative number of gene families shared by 100% of the genomes. The evolution of the average number of new genes per genome is also reported (in green). **(E)** Boxplots of the number of persistent (in gold), accessory (in grey) and singleton (in green) gene families per genome. When considering 1,294 genomes, there is on average 2,486 persistent proteins and only 26 singletons per genome.

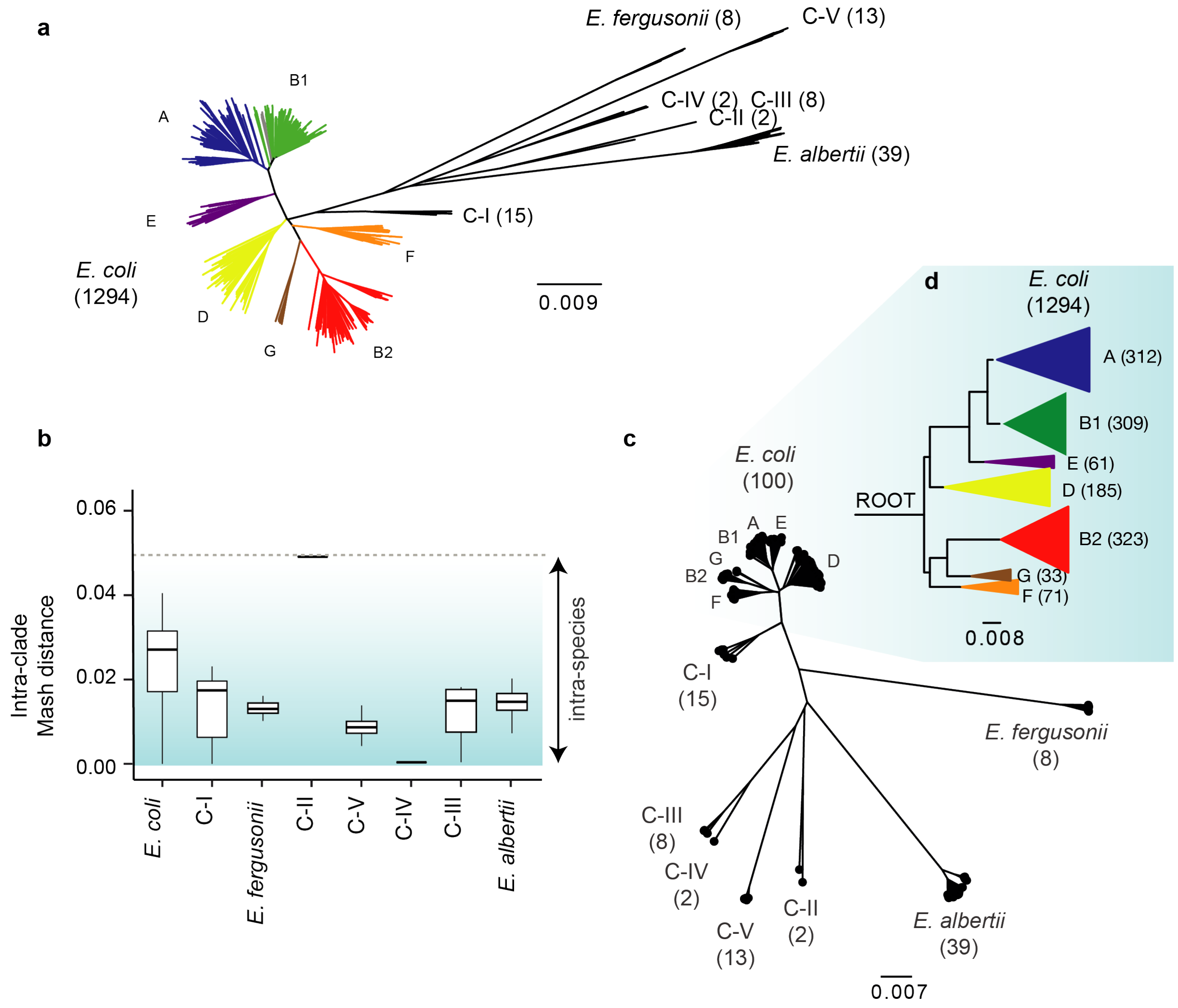


**Supplementary Figure 3: The genus and species phylogenetic trees. (a)** Distance tree of 1,294 Australian *E. coli* and 86 outgroups genomes performed from the matrix of mash distances computed between all pairs of genomes using bionj. The number of genomes in each species (or clade) was indicated. The different phylogroups of *E. coli* were displayed : A (in blue), B1 (in green), E (in purple), D (in yellow), F (in orange), G (in brown) and B2 (in red). **(b)** Boxplot of the mash distances computed between all pairs of genomes belonging to the same species (or clades). In both cases, the maximal mash distance was lower than 0.05. For *E. coli* species*,* the median was around 0.027 and the maximal value was 0.04. **(c)** Phylogenetic tree of 100 Australian *E. coli* genomes representative to the diversity of the dataset and 86 outgroups genomes performed from the persistent-genome of the genus with IQ-TREE under the GTR+F+I+G4 model. We made 1,000 ultra-fast boostrap to assess the robustness of the topology of the tree. We found that all boostrap supports were higher than 95%. **(d)** We rooted the species phylogenetic tree from the genus phylogenetic tree. The resulting rooted species tree was reported, and for simplicity, the main phylogenetic groups were collapsed.

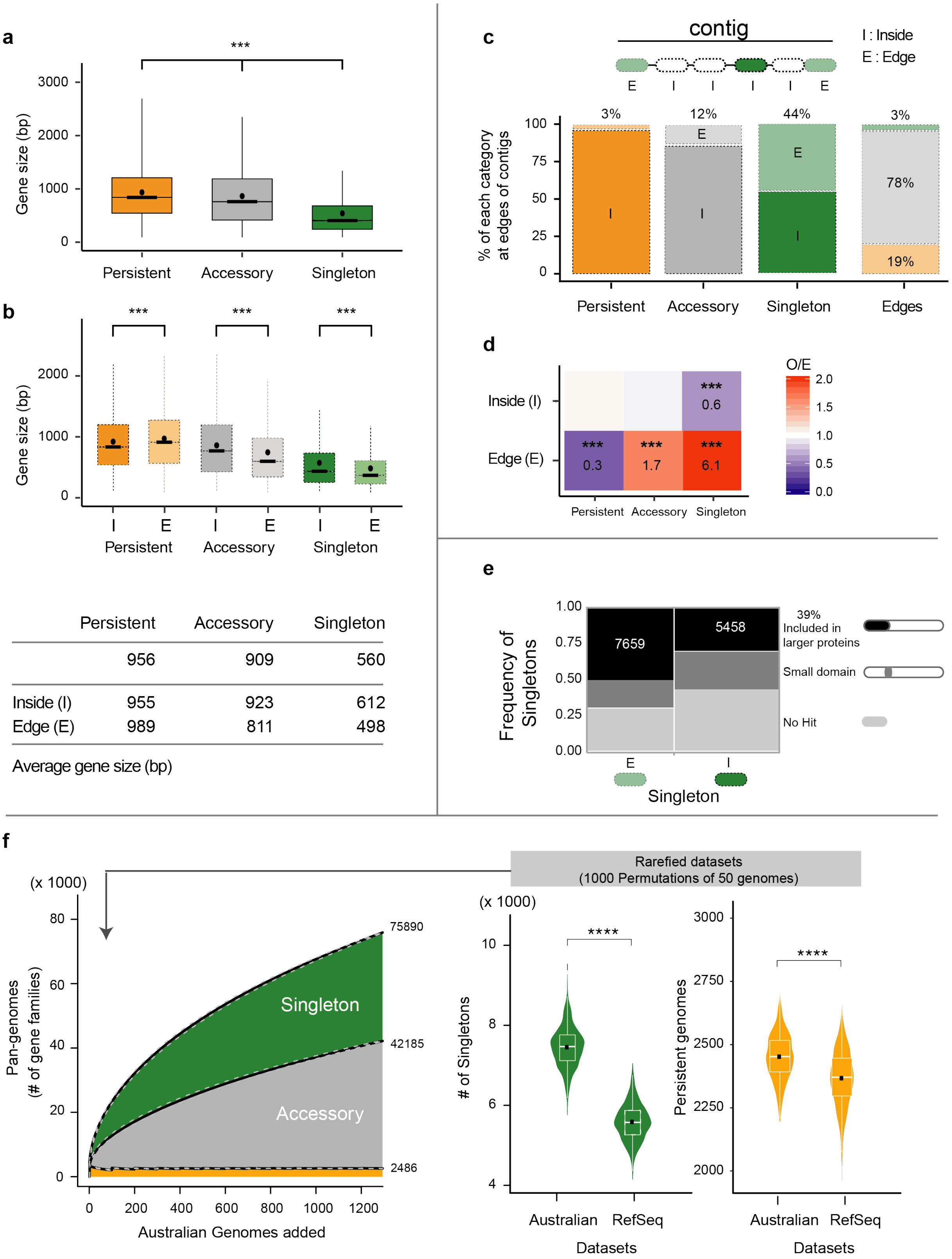


| | Persistent | Accessory | Singleton |
|---|---|---|---|
| | 956 | 909 | 560 |
| Inside (I) | 955 | 923 | 612 |
| Edge (E) | 989 | 811 | 498 |
| Average gene size (bp) | | | |

**Supplementary Figure 4: Singleton characterization. (a)** Boxplots of gene size (bp) in the three categories of gene families, *i.e.*, persistent (in gold), accessory (in grey) and singleton (in green). The average was represented by a black dot. The pairwise Wilcoxon Rank Sum test with bonferroni correction was applied to all comparisons (P<0.001 :***). **(b)** Same analysis as in **a**, but distinguishing the genomic location of the gene of each set : inside of contigs (I, dark color) or at the edge of contigs (E, light color). The average gene size for each case was reported in the table. (**c**) Percentage of genes located inside contigs (dark color) or at the edge of contigs (light color) in the 3 sets. The last column corresponds to the fraction of the 3 sets located at the edge of contigs. **(d)** Heatmap of the observed/expected (O/E) ratios of genes located inside or at the edges of contigs in the 3 sets. The ratio (O/E) was reported for all comparisons with a color code ranging from blue (under-representation) to red (over-representation). The level of significance of each Fisher's exact test was also indicated (P<0.001 :***). It was performed on each 2*2 contingency table. **(e)** Fraction of singletons with no hit (in light gey), with a small domain (in grey) or fully included (black) in larger accessory or persistent gene families (Text S2). **(f)** Violin plots of the number of singletons (in green) or persistent (in gold) observed in the rarefied Australian and RefSeq datasets. In each case, 1,000 permutations of 50 randomly selected genomes were performed (i.e., we used rarefied datasets). The boxplot is in white and the mean is represented by a black dot. While the average number of singletons is significantly higher (30% more) in the rarefied Australian dataset (Wilcoxon test, $P<10^{-4}$), the average number of persistent is also significantly higher (5% more, $P<10^{-4}$) than the rarefied RefSeq dataset. Singletons represent 43%, and 35% of the rarefied Australian and RefSeq pan-genomes, resp.

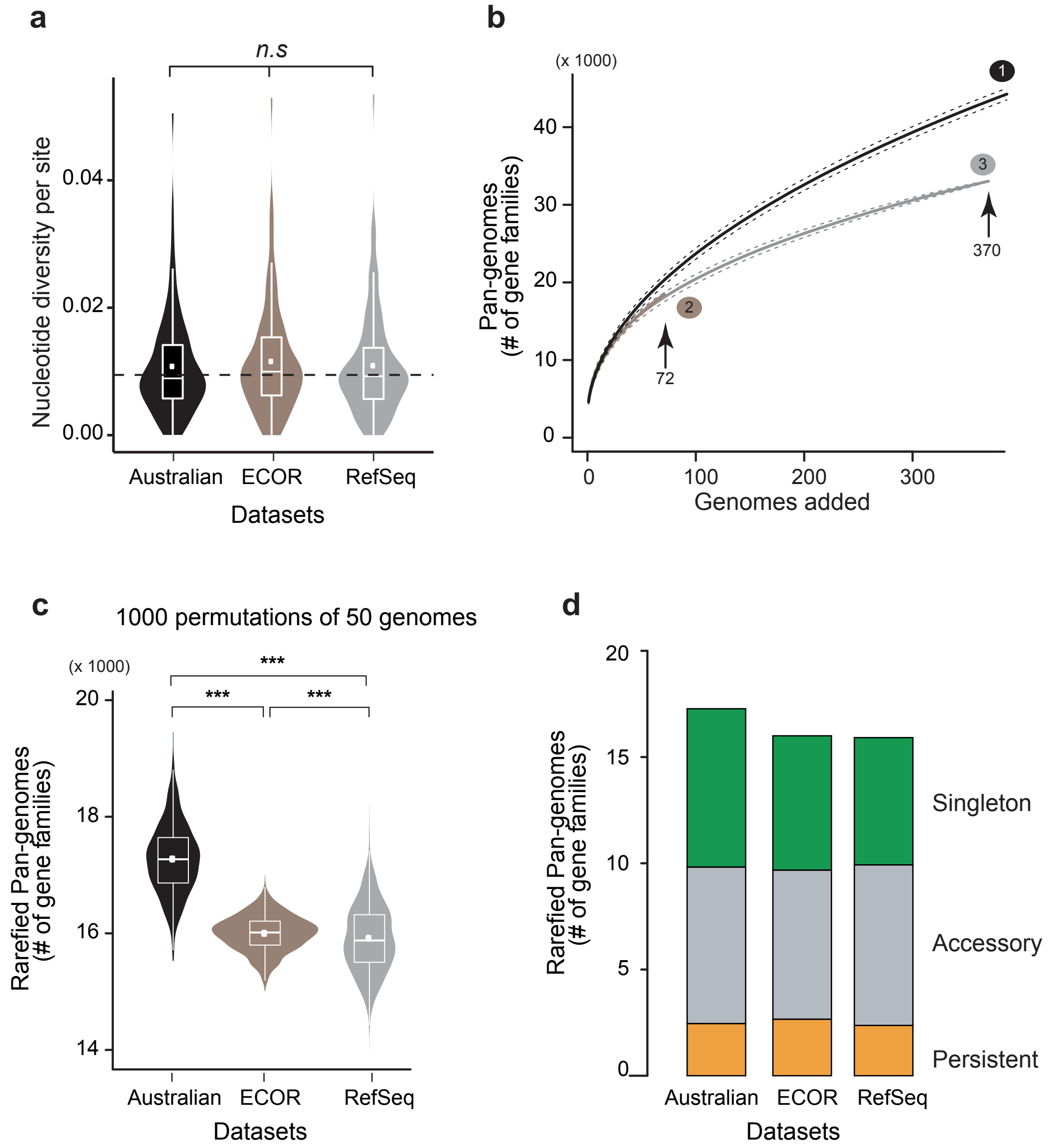


**Supplementary Figure 5: Comparison of Australian, ECOR and RefSeq datasets. (a)** Violin plots of the nucleotide diversity per site in the 3 datasets computed from the multiple alignments of 112 core gene families (see Methods). The pairwise Wilcoxon Rank Sum test with bonferroni correction was applied to all comparisons (P>0.05:ns). **(b)** Rarefaction curve of the full pan-genomes of the 3 datasets. In each case, we used 1,000 permutations (genomes orderings) and then averaged the results. **(c)** Violin plots of the size of the pan-genomes computed from the three rarefied datasets: In each case, 1,000 permutations of 50 randomly selected genomes were performed to calculated the rarefied pan-genomes. The pairwise Wilcoxon Rank Sum test with bonferroni correction was applied to all comparisons ($P<10^{-3}$ :***). **(d)** Average number of persistent (in gold), accessory (in grey) and singleton (in green) in the rarefied pan-genomes of each dataset.

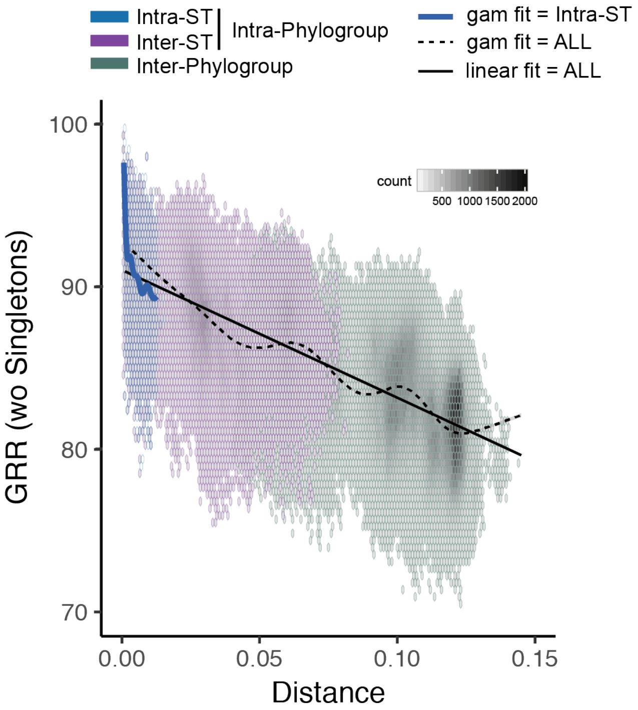


**Supplementary Figure 6: Association between GRR (%, Gene Repertoire Relatedness) and the patristic distance of each pair of genomes.** Here, the GRR were computed excluding singletons in all genomes. Due to the large amount of comparisons (points), we divided the plot area in regular hexagons. Color intensity is proportional to the number of cases (count) in each hexagon. The linear fit (full line, linear model (lm)) and the spline fit (dash line, generalized additive model (gam)) were reported for the whole (in black, all the species) or the intra-ST (in blue) comparisons. There was a significant negative correlation between GRR and the patristic distance (spearman's rho = -0.69, $P<10^{-4}$). The summary of the linear fit was : Y=90.722391-76.2919X, $R^2$=0.50,$P<10^{-4}$. Hence, with or without singletons, the results were similar.

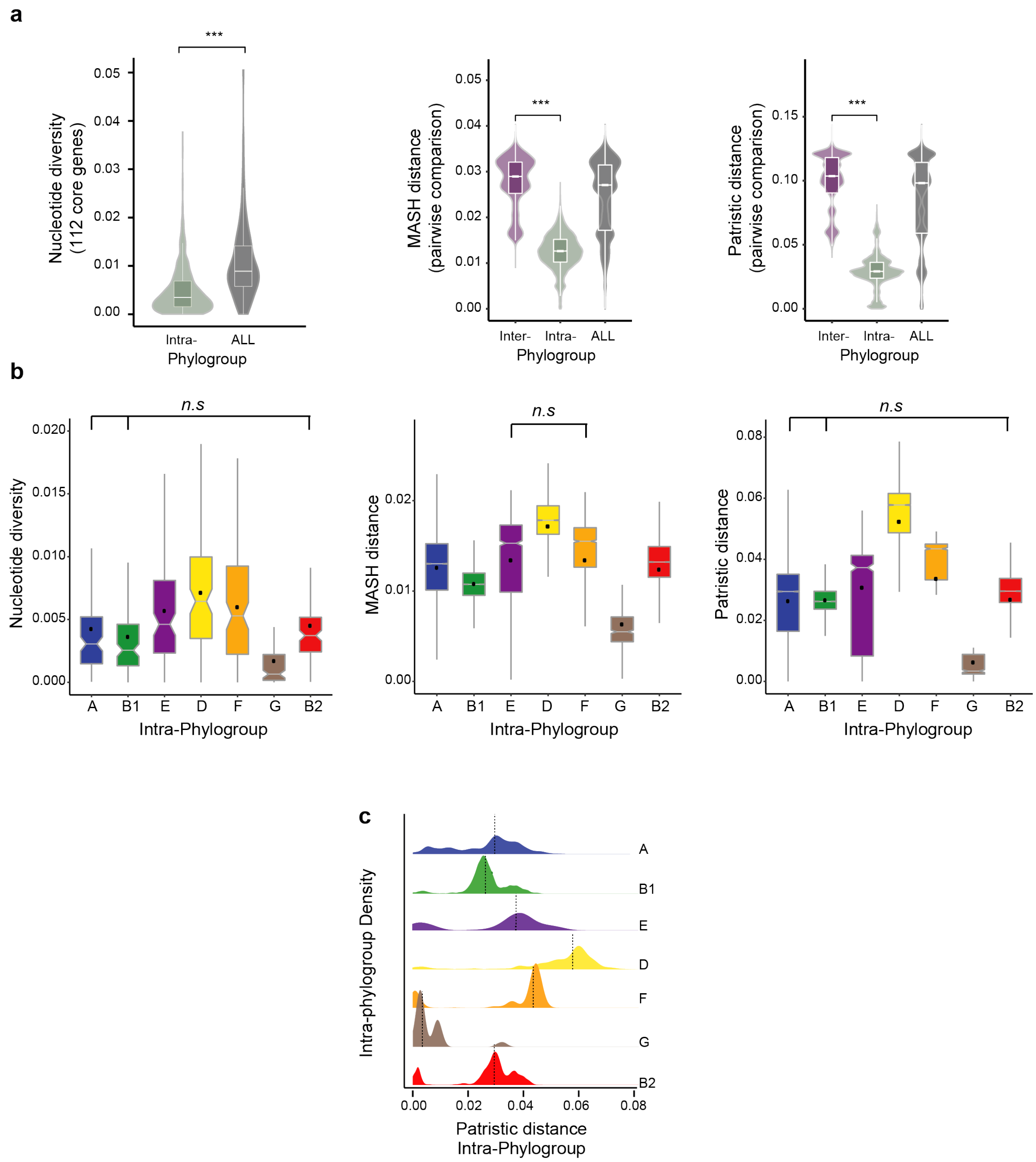


**Supplementary Figure 7: Intra- and Inter-phylogroup genetic diversity.** (**a**) Violin plots of the nucleotide diversity per site (left), the MASH (center) and the patristic distances (right) computed with/between genomes belonging to the same phylogroup (intra-phylogroup, in seagreen), to different phylogroups (inter-phylogroup, in purple), or all together (ALL, in darkgrey). In all cases, intra- and inter-phylogroup distributions were significantly different (Wilcoxon tests, $P<10^{-4}$). **(b)** Boxplots of the nucleotide diversity (left), the MASH (center) and the patristic distances (right) computed with/between genomes in each phylogroup. The pairwise Wilcoxon Rank Sum test with bonferroni correction was applied to all comparisons. Here, only the non-significant (ns : P>=0.05) comparisons were indicated, all other were higly significant $P<10^{-4}$. **(c)** Density of the patristic distances between all pairs of genomes of the same phylogroup (*intra-phylogroup*). The dash vertical line corresponds to the median of each distribution. (**a-b-c**) In all cases, similar results were obtained with rarefied datasets (i.e., comparing 50 randomly selected genomes in each groups, thus ignoring the small G phylogroup).

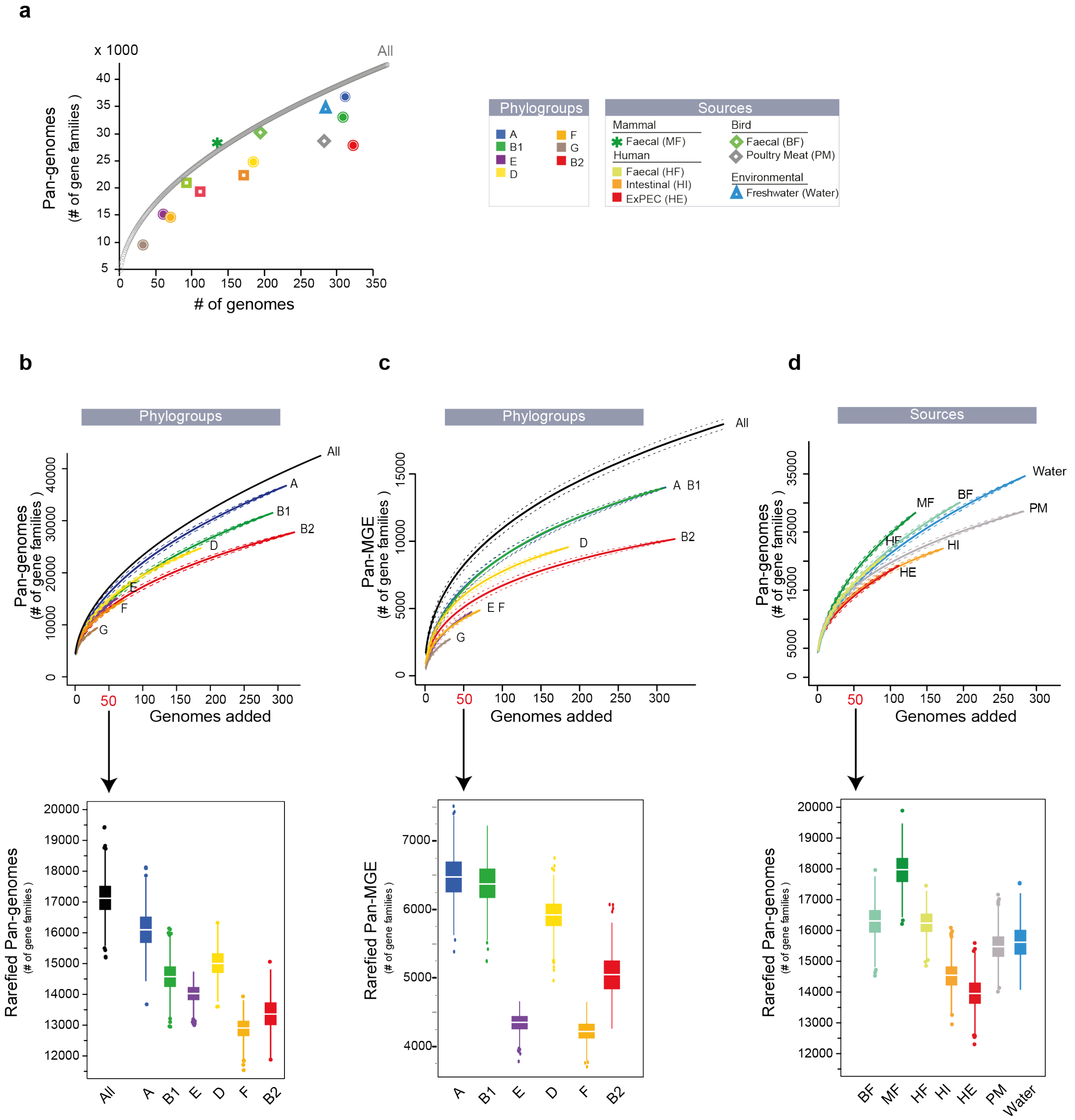


**Supplementary Figure 8: Pan-genomes, Pan-MGE, and rarefied Pan-genomes of each phylogroup and isolation source. (a)** Size of the pan-genome in each phylogroup and in each isolation source. The pan-genome sizes were correlated to the number of genomes in each group, even after excluding the singletons from the analysis (both, adjusted $R^2>0.88$, $P<10^{-4}$). The Rarefaction curve of the pan-genomes of the full dataset was also reported (All, in black). Rarefaction curves of **(b)** the pan-genomes of each phylogroup and of the full dataset (All), **(c)** the gene-families associated to MGE in each phylogroup and in the full dataset (All), **(d)** the pan-genomes of each isolation sources. In each case, (i) we used 1,000 permutations (genomes orderings) and then averaged the results (full line=mean, dash line=s.d), (ii) the pan-genomes remained open (with an *alpha* lower than one, see methods) that we considered them as a whole or without singletons, (iii) the boxplots of the rarefied pan-genomes (using a number of genomes = 50) were reported. The color code used was displayed in the insert (top right).

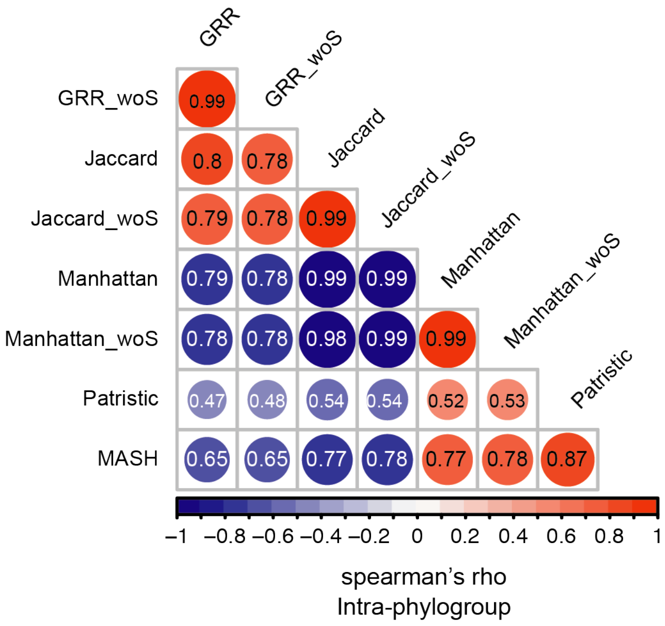


**Supplementary Figure 9: Correlation between the different distances and indexes**, *i.e.*, GRR, Manhattan, Jaccard,MASH and patristic, computed between pairs of genomes belonging to the same phylogroup (intra-phylogroup) with the whole dataset or excluding singletons (woS). Spearman's rank correlation rho matrix. Positive correlations were displayed in red and negative correlations in blue color. Color intensity and the size of the circle were proportional to the correlation coefficients. In the bottom side of the correlogram, the legend color shows the correlation coefficients and the corresponding colors. The p-value of each correlation was highly significant ($P<10^{-4}$). We found similar results with rarefied datasets, i.e., considering only 50 randomly selected genomes in each phylogroup. We also found higher correlation coefficients using all the comparisons (intra- and inter-phylogroup).

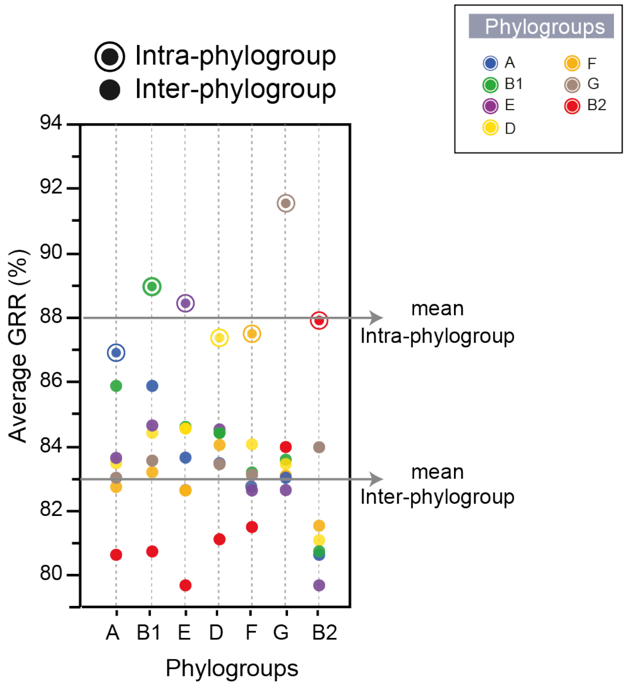


**Supplementary Figure 10: Gene repertoire relatedness (GRR) within and between phylogroups.** Average GRR (%) computed between pairs of genomes belonging to the same phylogroup (intra-phylogroup) and to different phylogroups (inter-phylogroup). The color code used was displayed in the insert (top right).

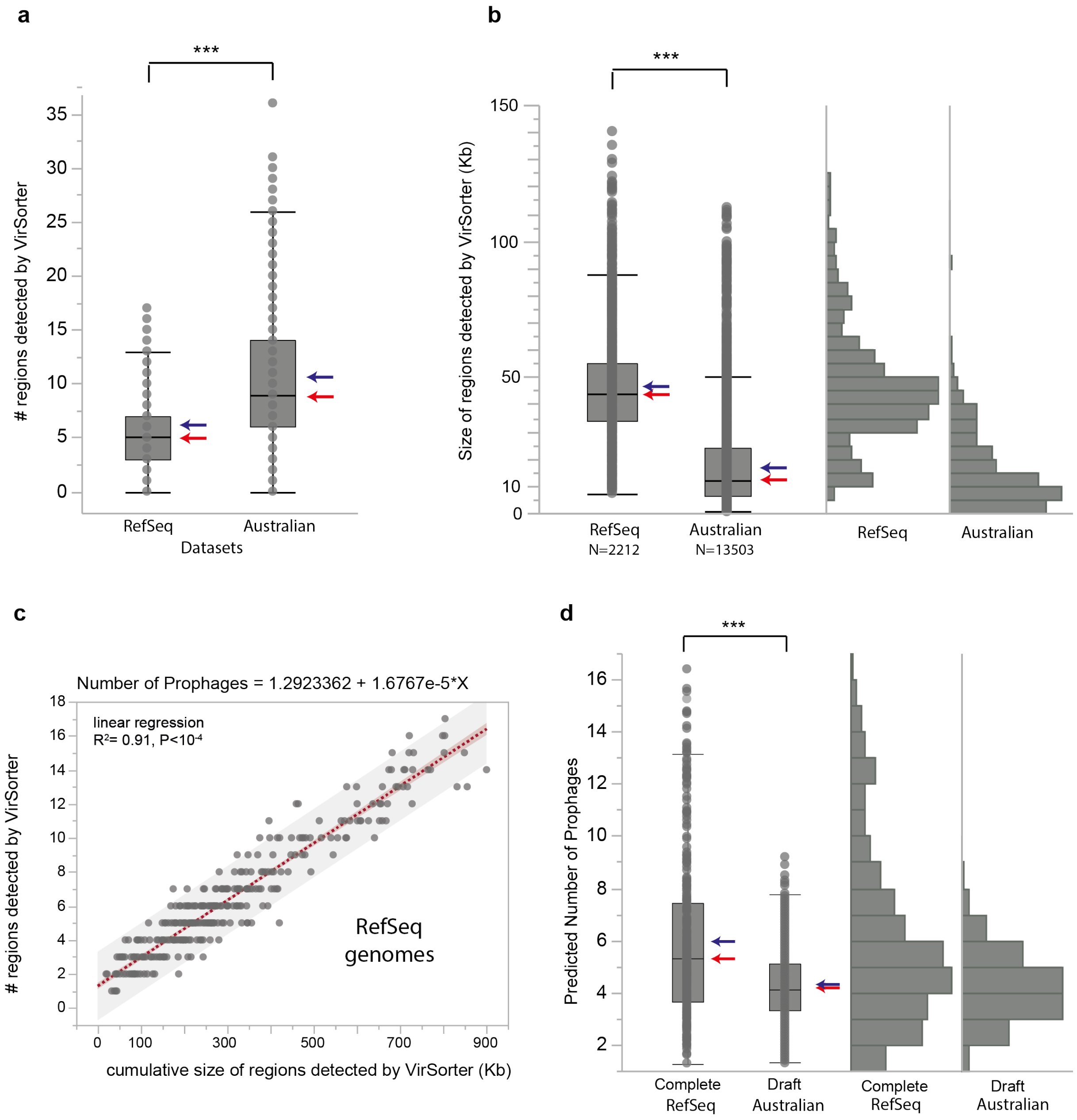


**Supplementary Figure 11: Detection and Estimation of the number of prophages. (a)** Boxplot of the number of regions detected as prophage-related by Virsorter in the 370 complete RefSeq GenBank genomes and in the 1,294 draft Australian genomes. These distributions were significantly different, on average the number of regions detected was significantly higher in draft than in complete genomes (Wilcoxon test, $P<10^{-4}$). **(b)** Boxplot and histogram of the size of the detected regions in complete and draft genomes. These distributions were significantly different (Wilcoxon test, $P<10^{-4}$). On average the regions were almost 4 times larger in the complete genomes than in draft genomes and few regions (644) in draft genomes had a typical size of known dsDNA phages (around 44kb). (**a-b**) showed that prophage elements were less assembled and were probably divided into several small contigs. The large regions (>60 kb) in complete genomes corresponded to tandem elements (consecutive on the genomic sequence). Thus, the number of detected regions did not correspond to the number of prophages either in the complete genomes (due to tandem elements) or in the drafts genomes (the elements being fragmented). **(c)** Strong association between the cumulative size of the detected regions (X) with the number of detected regions (Y). Linear regression (dash red line) and statistics were reported. **(d)**

Boxplot of the predicted number of prophage elements in both the complete and the draft genomes using the linear equation showed in (c) from the cumulative size of the regions detected by VirSorter. These distributions were significantly different (Wilcoxon test, $P<10^{-4}$). On average, there was 6.0 prophages in complete genomes, and 4.25 in draft genomes. The medians of the two data sets were closer reflecting probably the assembly problem related to the presence of prophages in tandem combined with the fact that they are often genetically close (most of them are lambdoids [13]). In each panel, the red arrow corresponds to the median and the blue arrow to the average of each distribution.

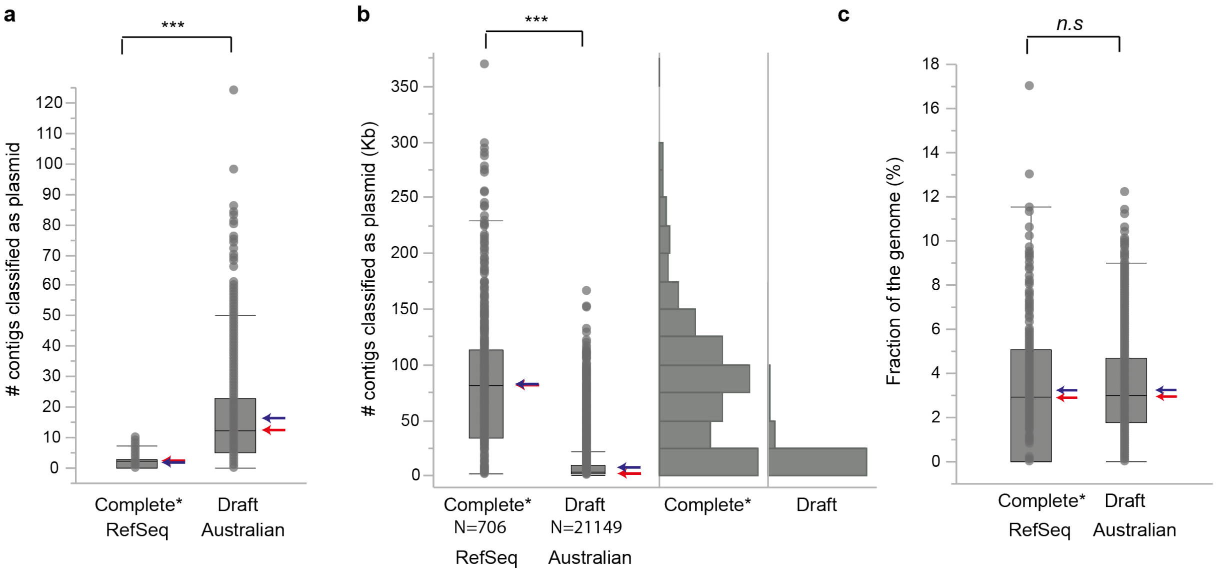


**Supplementary Figure 12: Detection of plasmid elements.** **(a)** Boxplot of the number of contigs classified as plasmid by PlaScope in the 370 complete RefSeq GenBank (Complete) genomes and in the 1294 draft Australian genomes (Draft). All the extrachromosomal replicons of the complete genomes were perfectly identified as plasmid elements by PlaScope. Hence, results based on the extrachromosomal replicons or on the contigs detected as plasmid by PlaScope were identical (Complete*). The average number of contigs was eight times larger in draft genomes than in complete genomes (15.4 vs 1.9) and reached up to 124 contigs. **(b)** Boxplot and histogram of the size of the contigs detected as plasmid in complete and draft genomes. These distributions were significantly different (Wilcoxon test, $P<10^{-4}$). On average the contigs were almost 10 times larger in the complete genomes than in draft genomes (81 kb vs. 8.9 kb). We identified 2347, 562 and 53 contigs larger than 20, 50 and 100 kb, resp. (**a-b**) showed that plasmid elements were poorly assembled and probably divided into several small contigs. **(c)** Boxplot of the fraction of the proteome encoding plasmid elements per genome (*i.e.*, the cumulative number of proteins located on contigs classified as plasmid divided by the total number of proteins of the genome) in complete and draft genomes. These distributions were similar (Wilcoxon test, $P>0.1$) with an average of 3.2% in both.

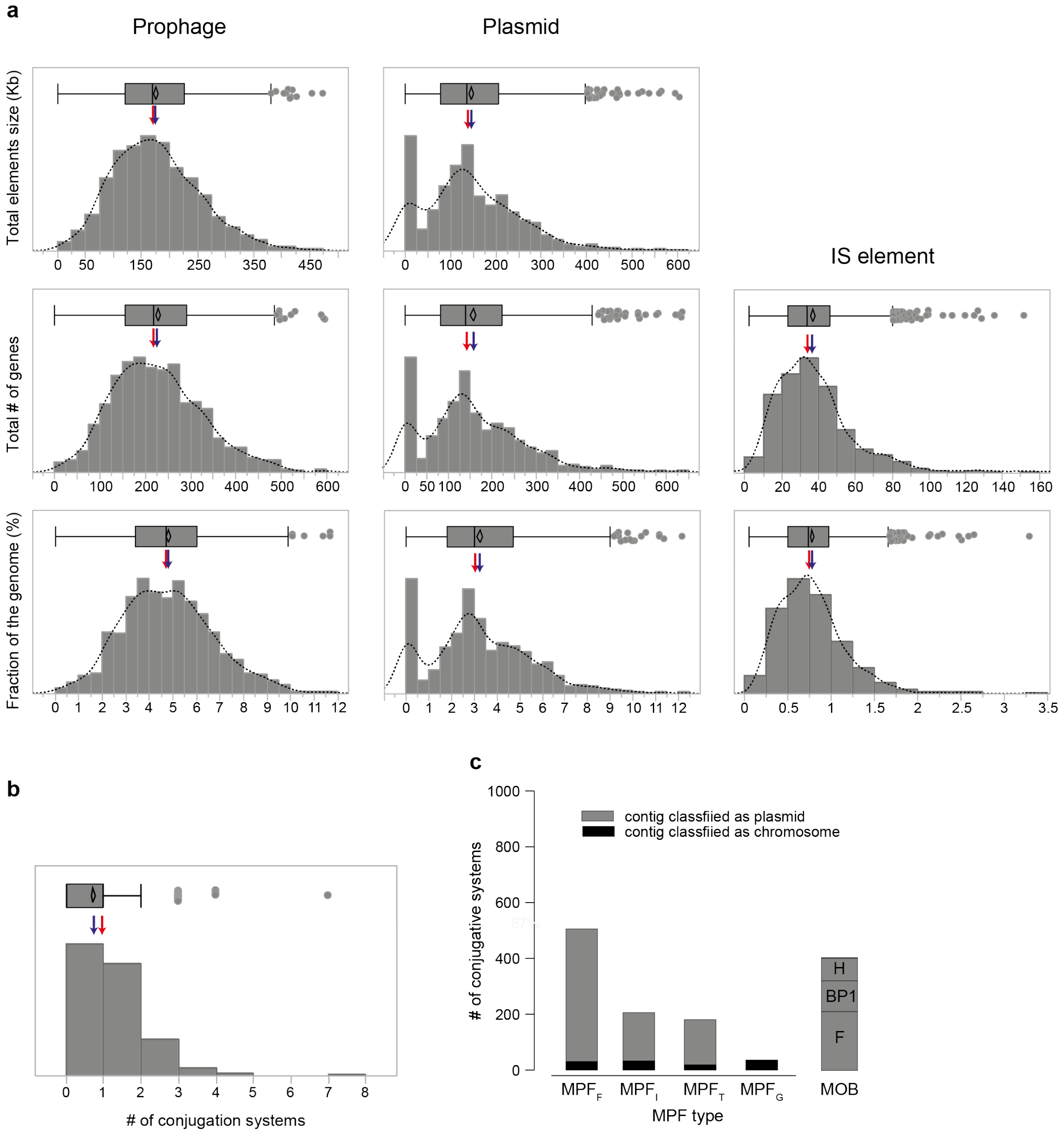


**Supplementary Figure 13: General genomic characteristics of the mobilome of Australian *E. coli*.** Three types of MGEs were detected, *i.e.*, prophage (left column), plasmid (middle columns) and IS elements (right column). **(a)** Histogram and boxplot of genomic features of each type of MGEs, *i.e*, the cumulative size of the elements per genome (Kb), the total number (#) of genes encoded by the elements per genome, the fraction of the genome encoding these elements per genome. For each case, the dash line corresponds to the smoothed curve, the red arrow to the median and the blue arrow to the average of each distribution. **(b)** Histogram and boxplot of the number of conjugation systems per genome. **(c)** Number of conjugative systems: (MPF) and isolated relaxases (MOB) detected in our dataset. The different MPF types were indicated and also their genomic location, *i.e.,* located on a contig classified as plasmid or as chromosome by PlaScope [14].

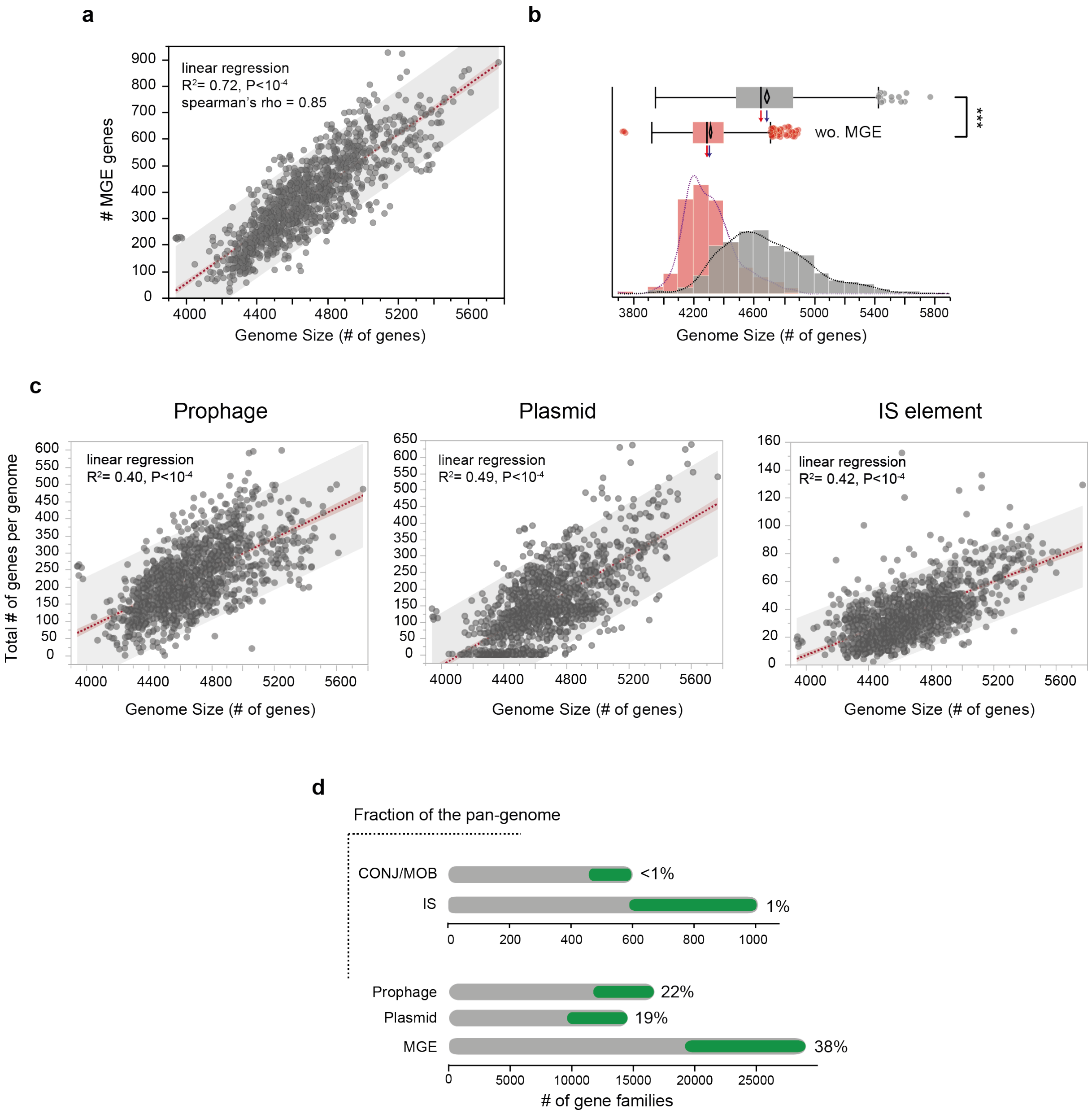


**Supplementary Figure 14: Contribution of MGEs to genome size variation. (a)** Association between the genome size (*i.e.*, # of genes per genome) and the total number of genes associated to the MGE elements. **(b)** Histogram and boxplot of the genome size (in grey), and of the genome size without MGE (in red), i.e., after removing all the genes encoding MGE elements (in red). These distributions were significantly different (Wilcoxon test, $P<10^{-4}$). **(c)** Same representation as in (a), but distinguishing the different types of MGEs, i.e., prophage, plasmid and IS elements. **(a-c)** We found a strong correlation in each case. Linear regression (dash red line) and statistics were reported. Similar results were obtained with the genome size (Mb). **(d)** Number of singletons (in green) and accessory gene families encoding MGEs. The fraction of the pan-genome encoding such elements was reported in each case (%).

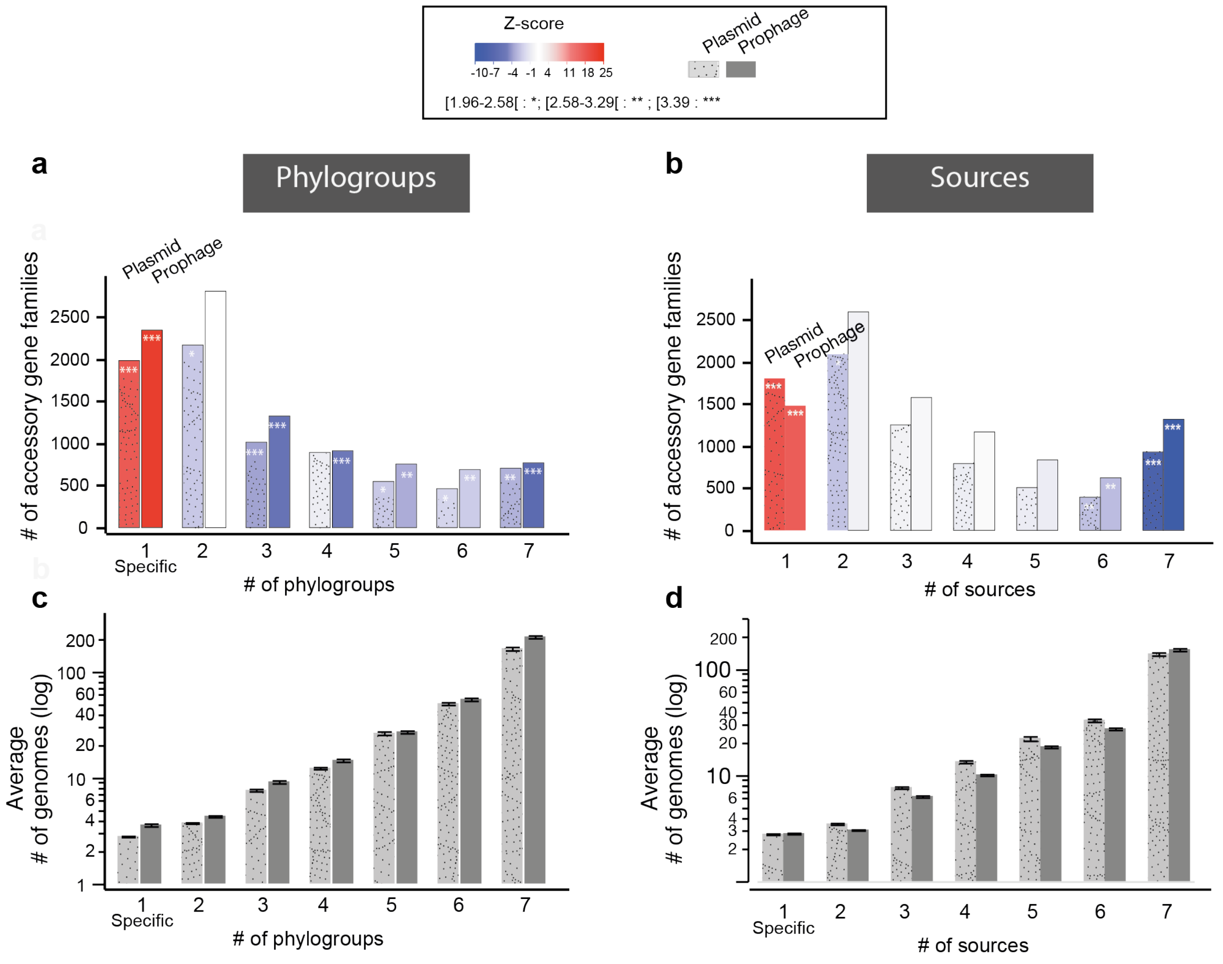


**Supplementary Figure 15: Distribution of gene families related to MGEs across phylogroups and sources.** Number of accessory gene families associated to prophage and plasmid present in one (i.e., phylogroup-specific) to seven phylogroups **(a),** or in one (i.e., source specific) to seven sources **(b)**. The Z-score obtained for the observed number with respect to the expected distribution (as in Fig. 3d, we randomized 1,000 times, only the phylogroup (a) or the source (b) assignment of genomes) was reported for each case with a color code ranging from blue (under-representation) to red (over-representation). The frequency of these families (average number of genomes) was also indicated in (c) for phylogroups, and in (d) for sources.

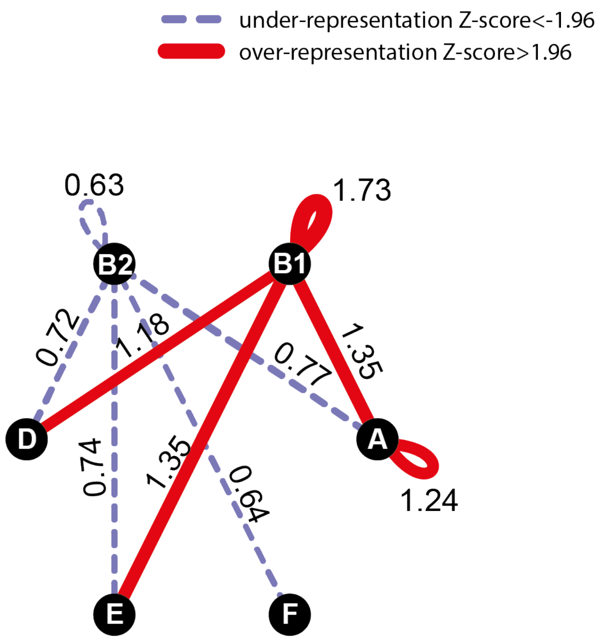


**Supplementary Figure 16: Network of recent co-occurence of gains (co-gains) of MGE genes within and between phylogroups.** Nodes are phylogroups and edges the O/E ratio of the number of pairs of MGE genes (from the same gene family) acquired in the terminal branches of the tree. Only significant O/E values (and edges) are plottted (|Z-score|>1.96). Under-represented values are in dash blue and over-represented in red (see Methods).

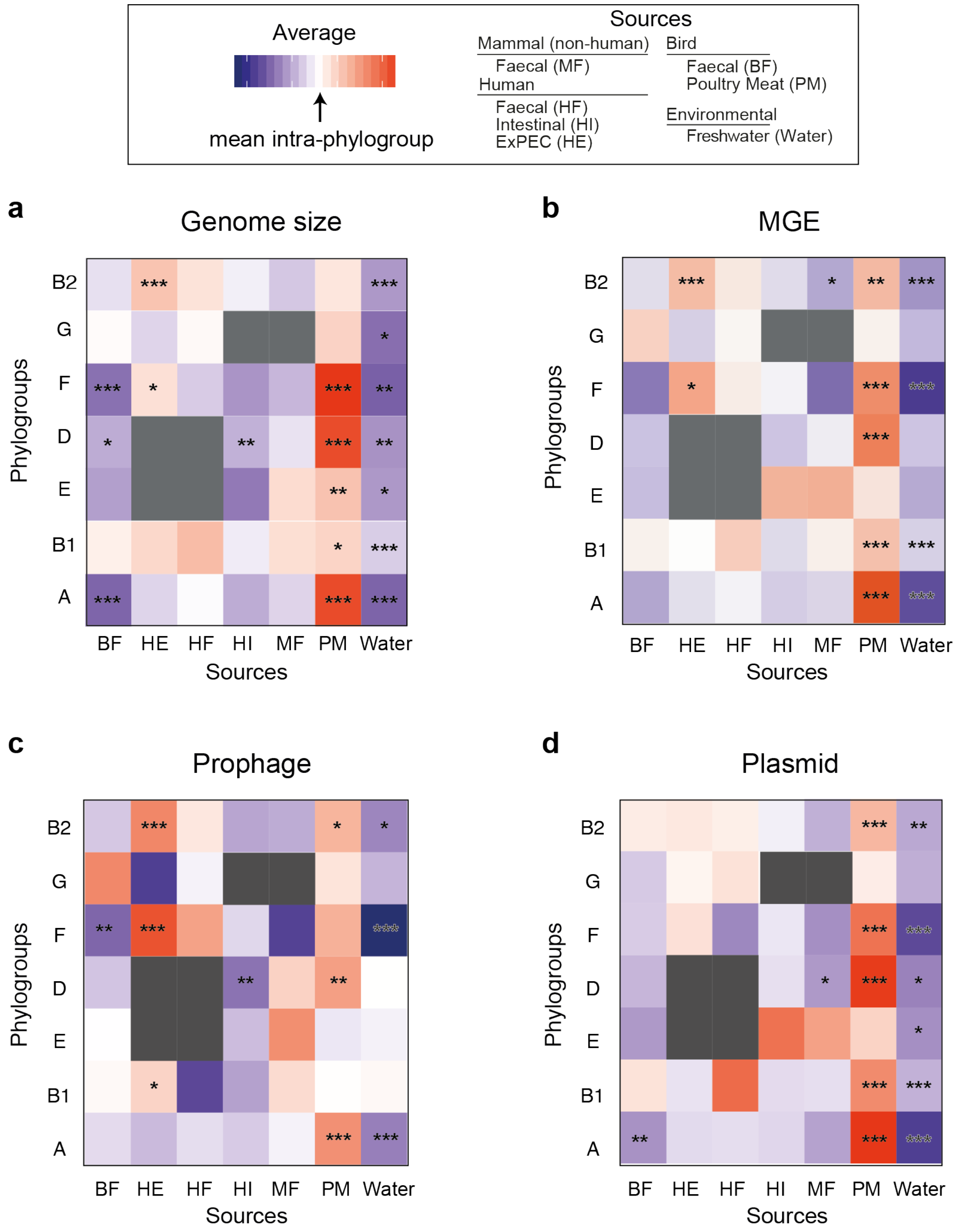


**Supplementary Figure 17: Genome size and MGE content according to sources within each phylogroup. (a)** Heatmap of the average genome size of strains from different sources in each phylogroup. The deviation to the overall intra-phylogroup mean (i.e., the average genome size of all strains belonging to a given phylogroup) was reported for all comparisons with a color code ranging from blue (below average) to red (above average). The level of significance of each ANOM test was indicated (P>=0.05 : ns ; P<0.05 : * ; P<0.01 : ** ; P<0.001 :***). It was performed within each phylogroup (each line). **(b-c-d)** Same representation as in (a), but in relation with the average number of genes associated to MGEs **(b)**, to prophage **(c)**, or plasmid elements **(d)**.

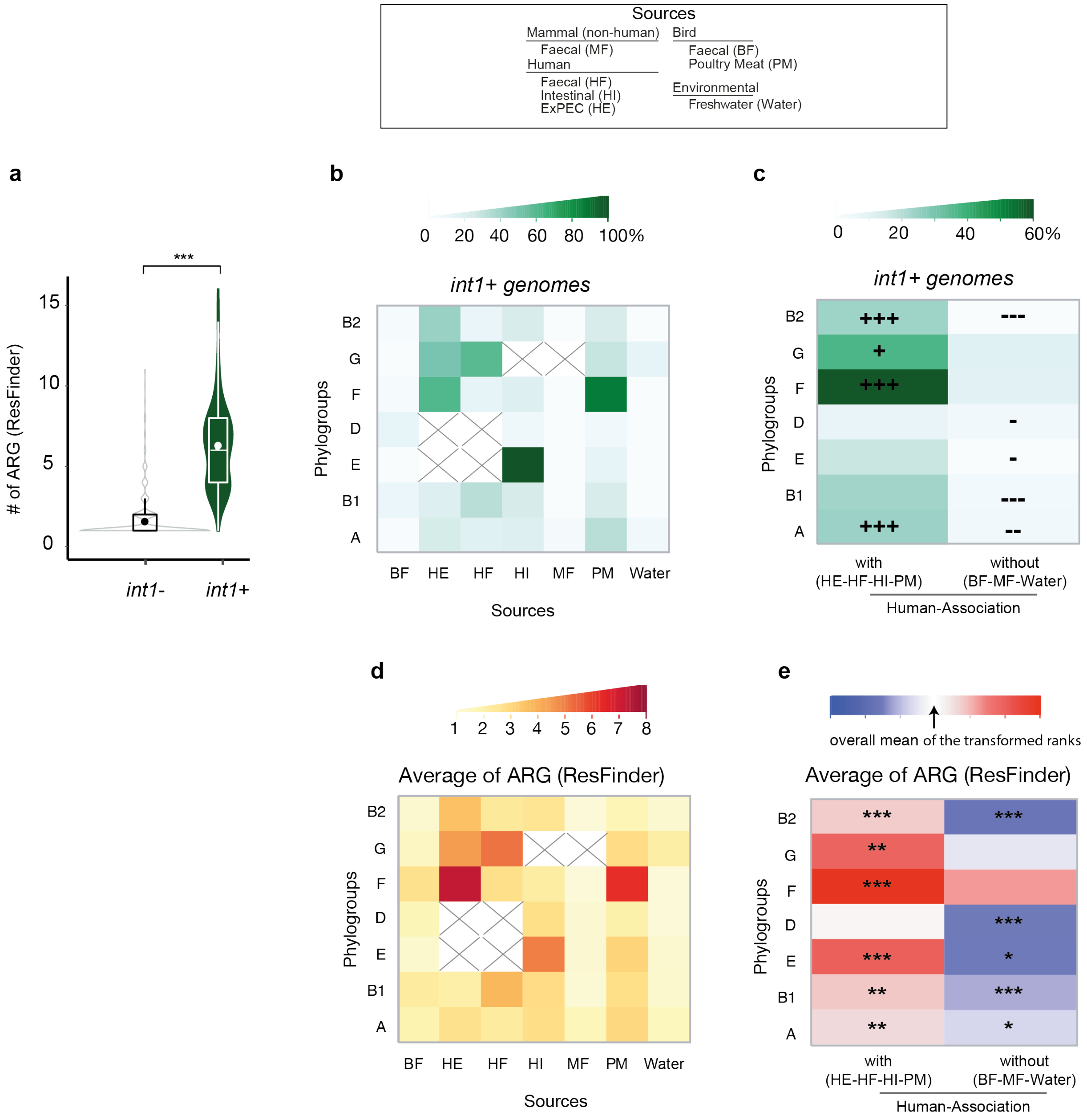


**Supplementary Figure 18: Association of integrons and ARGs with human (or domesticated animals). (a)** Violin plots of the number of ARGs in genomes encoding integron-integrase (*int1*+) or not (*int1*-). The level of significance of the Wilcoxon test was indicated ($P<10^{-3}$). **(b)** Heatmap of the proportion of genomes *int1*+ in each phylogroup and source. A cross marks the absence of data. **(c)** Same as in (b) but we merged sources related to human activity (with), or not directly associated to human (without). The level of significance of each ANOM for proportions test was indicated (P>=0.05 : ns ; P<0.05 : * ; P<0.01 : ** ; P<0.001 :***). Here, we compared response proportions for the X levels to the overall response proportion from the contingency table. This method uses the normal approximation to the binomial. Therefore, in some cases sample sizes were too small to be tested. **(d)** Heatmap of the average number of ARGs per genome in each phylogroup and source. **(e)** Heatmap of the average number of ARGs when we merged sources related (with) or not (without) to human activity. The level of significance of each non-parametric ANOM test (ANOM with Transformed Ranks) was indicated (P>=0.05 : ns ; P<0.05 : * ; P<0.01 : ** ; P<0.001 :***). The deviation to the overall mean (i.e., in all genomes) was reported for all comparisons with a color code ranging from blue (below average) to red (above average). The color code used was displayed in the top of each panel.

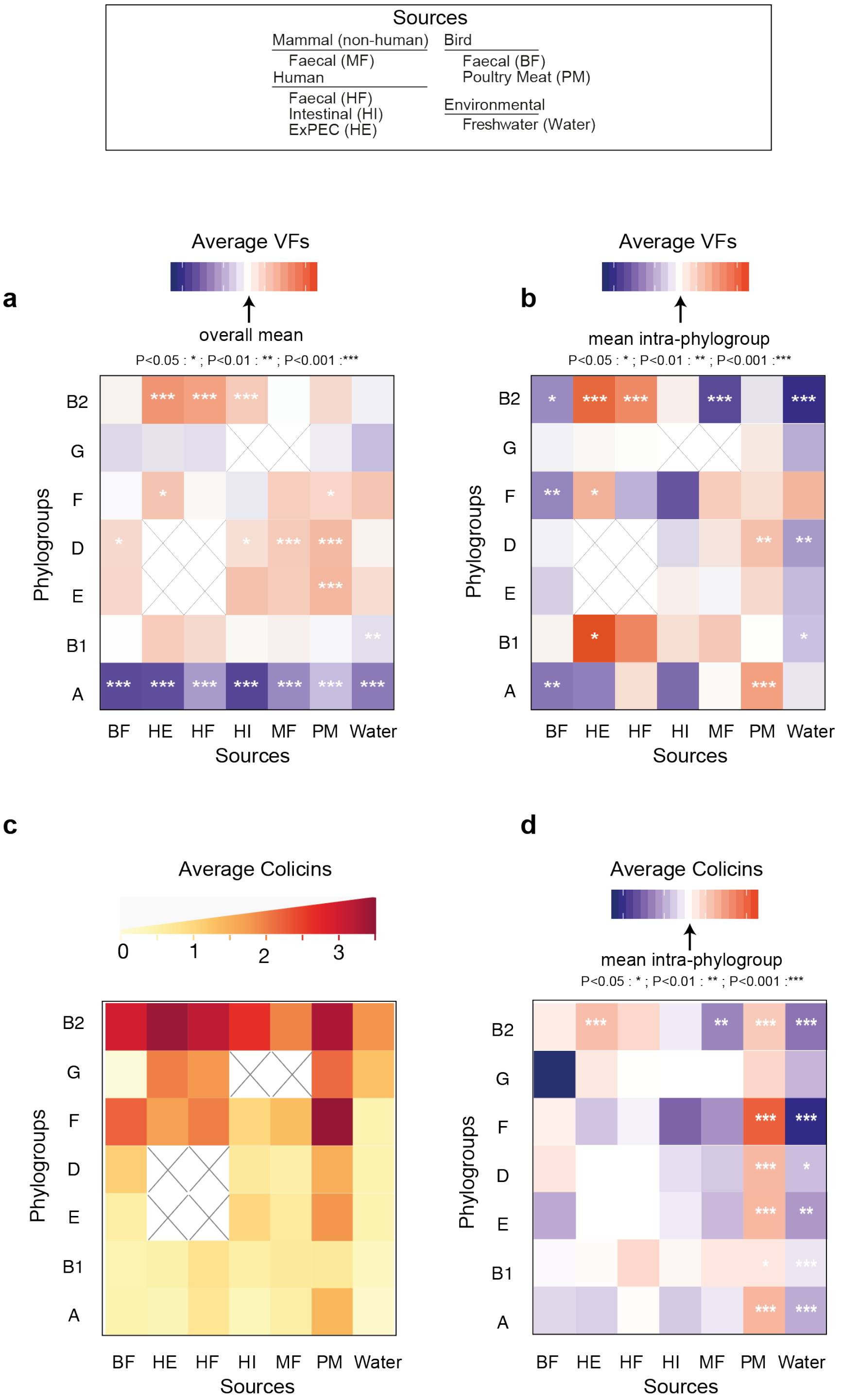


**Supplementary Figure 19: Distribution of VFs and Colicins MGEs across phylogroups and sources. (a-b)** Heatmap of the average number of VFs per strain from different sources in each phylogroup. The deviation to the overall mean (i.e., whole dataset, in **a**) or to the intra-phylogroup mean (i.e., the average number of all strains belonging to a given phylogroup, in **b**) was reported for all comparisons with a color code ranging from blue (below average) to red (above average). The level of significance of each ANOM test was indicated (P>=0.05 : ns ; P<0.05 : * ; P<0.01 : ** ; P<0.001 :***). It was performed within each phylogroup (each line, in **b**). **(c)** Heatmap of the average number of Colicins per genome in each phylogroup and source. **(d)** Same representation as in (b), but in relation with the average number of Colicins per genome.
